# The in-tissue molecular architecture of β-amyloid in the mammalian brain

**DOI:** 10.1101/2022.11.08.515609

**Authors:** Conny Leistner, Martin Wilkinson, Ailidh Burgess, Stanley Goodbody, Yong Xu, Susan Deuchars, Sheena E. Radford, Neil A. Ranson, René A. W. Frank

## Abstract

Amyloid plaques composed of extracellular focal deposition of Aβ fibrils are a hallmark of Alzheimer’s disease (AD). Cryo-EM structures of Aβ fibrils purified from human AD brain tissue post mortem have recently been determined. However, the molecular architecture of amyloid plaques in the context of fresh, unfixed mammalian brain tissue is unknown. Here, using cryogenic correlated light and electron tomography we report the native, *in situ* molecular architecture of Aβ fibrils in the brain of a mouse model containing the Arctic familial AD mutation (*App*^*NL-G-F*^) and an atomic model of Arctic Aβ fibril purified from the brains of these animals. We show that in-tissue Aβ fibrils are arranged in a lattice or in parallel bundles within a plaque, and are interdigitated by subcellular compartments, exosomes, extracellular droplets and extracellular multilamellar bodies. At the atomic level, the Arctic Aβ fibril differs significantly from earlier structures of Aβ amyloid extracted from *App*^*NL-F*^ mice models and human AD brain tissue, showing a striking effect of the Arctic mutation (E22G) on fibril structure. Cryo-electron tomography of *ex vivo* purified and in-tissue amyloid revealed an ensemble of additional fibrillar species, including thin protofilament-like rods and branched fibrils. Together, these results provide a structural model for the dense network architecture that characterises β-amyloid plaque pathology.

## INTRODUCTION

Alzheimer’s disease (AD) results in cognitive decline and brain atrophy that is characterized by multiple pathologies, including the formation of abnormal extracellular protein deposits of β-amyloid (Aβ), intracellular tangles of tau, alongside neuroinflammation and the loss of neurons and synapses^1^. Mutations identified in familial forms of AD (FAD) indicate a causal role for the amyloid precursor protein (APP) and presenilin (PSEN1/PSEN2) genes, which encode the precursor of Aβ and the γ-secretase enzyme that catalyses the final step of Aβ peptide production, respectively^2,3^. γ-Secretase produces Aβ peptides that vary in length, of which Aβ_1-40_ and Aβ_1-42_ are most abundant. These peptides are highly aggregation prone, assembling into diffusible, low-molecular weight oligomers or protofibrils that precede the formation of larger Aβ fibrils^4^. Over decades, Aβ peptides, particularly Aβ_1-42_ ^5^, accumulate and form amyloid plaques in the parenchyma of the AD brain^2^. Amyloid plaques have been categorised as diffuse, dense-cored, fibrillar or neuritic, all of which contain fibrillar Aβ deposits^6^. Conventional EM of plastic embedded tissue from post mortem AD brain^7,8^ and FAD animal models^9^ suggest that plaques are composed of parallel bundles and a lattice of Aβ fibrils. Additionally, conformation-specific fluorescent dyes suggest a heterogeneity of Aβ conformations within distinct regions of amyloid plaques^10,11^.

Structures of amyloid fibrils of Aβ have been determined recently by cryoEM studies of fibrils assembled *in vitro* from recombinant or synthetic Aβ_1-42_ 12, as well *ex vivo* fibrils purified from post mortem human AD brain and FAD mouse models^13,14^. These *ex vivo* samples yielded two structural arrangements for fibrils within Aβ_1-42_ amyloid, with type I fibrils associated with sporadic disease and type II associated with familial disease and other brain pathologies^13^. Both forms differ from Aβ_1-42_ fibrils prepared *in vitro*^12^ and from Aβ_1-40_ prepared from the meninges of cerebral amyloid angiopathy (CAA) cases^15^. However, the molecular architecture and organization of β-amyloid within plaques of fresh, unfixed, brain tissue remained unknown.

Here we sought to visualize Aβ fibrillar plaques *in situ* using a homozygous knockin FAD mouse model (*App*^*NL-G-F/NL-G-F*^)^16^ by preparing fresh, hydrated vitrified tissue samples for cryogenic correlated light and electron microscopy (cryoCLEM) and cryo-electron tomography (cryoET). We also determined the *ex vivo* near-atomic resolution structure of Aβ fibrils purified from the same *App*^*NL-G-F*^ mouse brains by single particle cryo-EM, which revealed a dramatic change in Aβ fibril structure caused by the Arctic mutation (E22G). Analysis of the fibrils within in-tissue tomograms revealed the presence of fully assembled fibrils, along with protofilament-like rods which may describe early assembly intermediates, and branched fibrils, suggestive of secondary nucleation mechanisms occurring *in vivo*. These data describe a dense network of fibrils, interdigitated with non-amyloid constituents that defines the in-tissue 3D molecular architecture of the amyloid plaque.

## RESULTS

To determine the molecular architecture of pathology within fresh, intact tissue we developed a cryoCLEM workflow that we applied to 11-14 month-old *App*^*NL-G-F*^ mice^16^. These animals have a humanized mouse *App* gene containing three familial Alzheimer’s disease mutations (Swedish, Beyruthian, and Arctic)^16^. The Swedish and Beyruthian mutations are located upstream and downstream of the coding region for the Aβ peptide, and increase the overall Aβ concentration, and the ratio of Aβ_1-42_:Aβ_1-40_, respectively. In contrast, the Artic mutation is situated within the Aβ peptide coding sequence (App E693G, Aβ E22G) and is thought to increase the generation of Aβ protofibrils^17^. *App*^*NL-G-F*^ mice develop β-amyloid plaques, neuroinflammation, damaged synapses, and behavioural phenotypes, without ectopic over-expression of APP^18^. To identify amyloid pathology within fresh tissue, mice received an intraperitoneal injection of a fluorescent amyloid dye, methoxy-X04 (MX04; **Fig. 1a**)^19^. Immunohistochemical fluorescence imaging confirmed MX04 detected β-amyloid in *App*^*NL-G-F*^ mice (**Extended Data Fig. 1a,b**).

**Figure 1.**
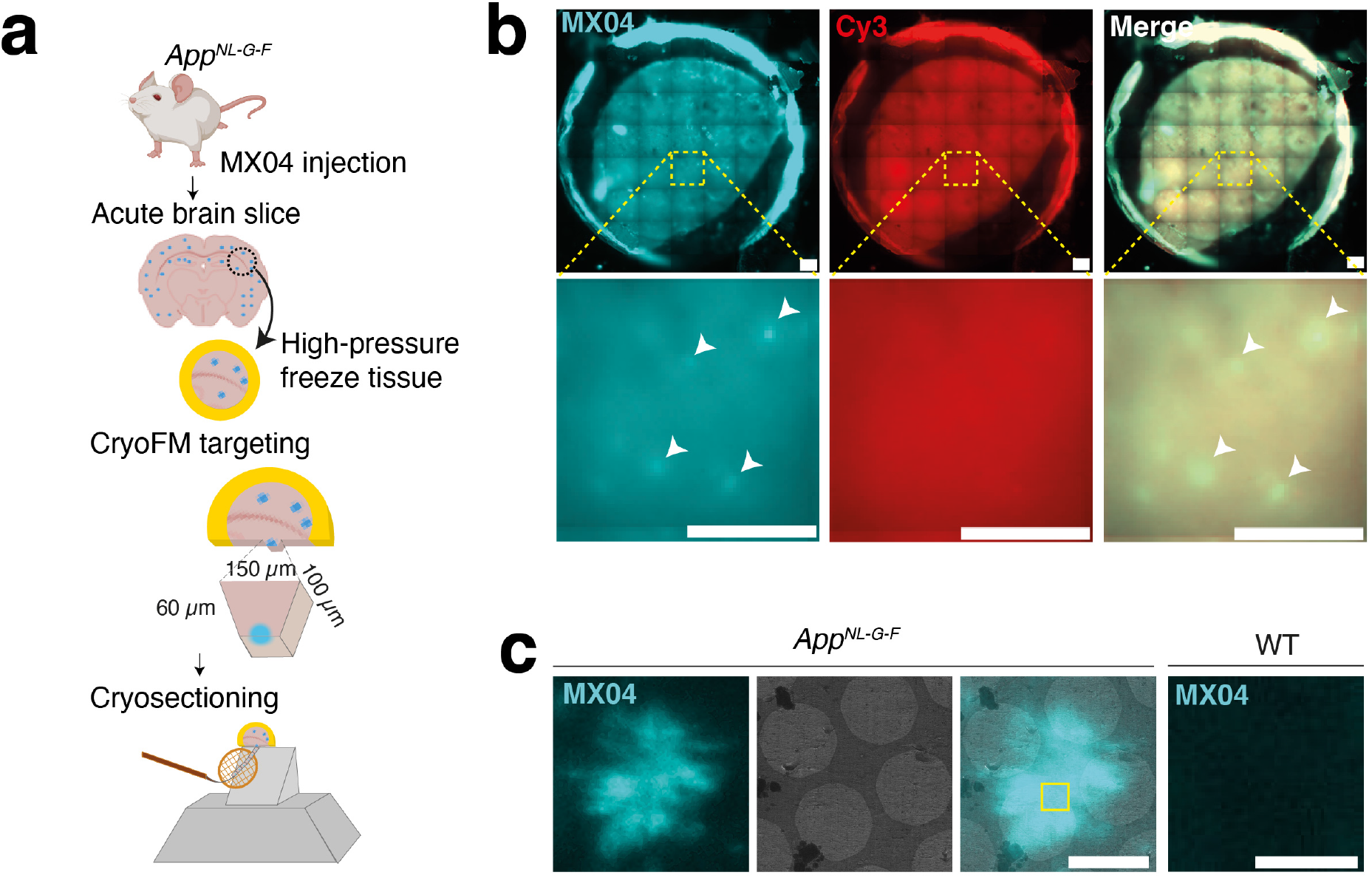
Correlated light and electron microscopy to target the in-tissue pathology in the mammalian brain. **(a)** Schematic showing correlative imaging work-flow to prepare fresh tissue for in-tissue cryo-electron tomography. *App*^*NL-G-F*^ knockin mice were systemically administered with the amyloid fluorescent dye (methoxy-X04, MX04). Fresh acute brain slices were prepared, and tissue biopsies were high-pressure frozen. MX04-labelled plaques were detected by cryogenic fluorescence microscopy (cryoFM) within vitreous frozen tissue and used to target trimming of a trapezoid tissue stub to contain an amyloid plaque by cryo-ultramicrotomy. 70-150 nm thick tissue CEMOVIS/cryo-sections were collected from the front face of the amyloid plaque-containing stub and attached to the holy carbon support of an EM grid. Cryo-sections were 70-150 nm thick and encompassed an area of ∼130 x ∼90 μm **(b)** CryoFM imaging of a high pressure frozen *App*^*NL-G-F*^ tissue biopsy. *Top panels*, image of whole tissue biopsy within circular gold carriers, and *bottom panels*, close-up. *Left panels*, Detection of MX04-labelled amyloid plaques (cyan puncta). *Middle panels*, Control image of samples stained with Cy3 (excitation and emission of 550 and 620 nm, respectively) to detect non-specific autofluorescence and contamination. *Right panels*, Merged image. Arrowheads indicate prominent amyloid plaques. Scale bars, 200 μm. **(c)** Cryogenic correlated light and electron microscopy (cryoCLEM) to locate amyloid within cryo-sections of *App*^*NL-G-F*^ tissue and wild-type tissue control. *First left panel*, cryoFM detection of MX04-labelled amyloid plaque (cyan). *Second left panel*, cryoEM image of amyloid amyloid-containing tissue cryo-section. *Second right panel*, merged imaging showing aligned cryoFM and cryoEM image. Yellow box, region of amyloid plaque selected to collect the in-tissue tomogram shown in **Fig. 2**. *First right panel*, MX04-labelled wild-type tissue section serving as a control for the detection of amyloid by MX04. Scale bars, 4 μm.

To cryopreserve anatomically intact tissue, 2 mm cortical biopsies of acute brain slices were high-pressure frozen and imaged by cryogenic fluorescence microscopy (cryoFM), identifying the locations of MX04-labelled amyloid plaques (**Fig. 1a,b**). We used these cryoFM maps of amyloid deposits to target the preparation of tissue cryo-sections by trimming a 60 x ∼100 x ∼150 μm stub of tissue containing a single MX04-labelled amyloid plaque, from which 70-150 nm thick tissue cryo-sections were collected and attached to an EM grid support (**Fig. 1a**)^20–22^. CryoFM of *App*^*NL-G-F*^ tissue cryo-sections revealed 5-20 μm star-shaped or round β-amyloid pathology (**Fig. 1c**). These were similar to plaques observed by fluorescence imaging of fixed samples and were absent in MX04-injected wild-type control samples (**Fig. 1c and Extended Data Fig. 1b,c**). MX04-labelled amyloid plaques were mapped on medium magnification electron micrographs by cryoCLEM (**Fig. 1c**) to target areas for the collection of tomographic tilt series, which each encompassed a 1.3 μm^2^ area of the tissue cryosection. We collected 23 tomograms (4 with and 19 without a Volta phase plate) sampling central and peripheral regions of MX04-labelled plaques (**Extended Data Table 1**).

### The native 3D molecular architecture of amyloid plaques

Reconstructing 3D cryotomographic volumes revealed the native, in-tissue cellular and molecular architecture of amyloid pathology (**Fig. 2**) (see **Methods** describing criteria for identifying macromolecular and cellular constituents within cryoET data). Fibrils were present within all tomographic volumes that spatially correlated with the MX04 fluorescence by cryoCLEM (**Fig. 2a-c and Fig. 3**). The results revealed a dense array of fibrils that comprise the amyloid plaques in which the fibrils are arranged as a lattice or in parallel bundles (**Extended Data Fig. 2**), consistent with previous observations using conventional (fixed, plastic embedded, heavy metal-stained) EM^7–9,23^.

**Figure 2.**
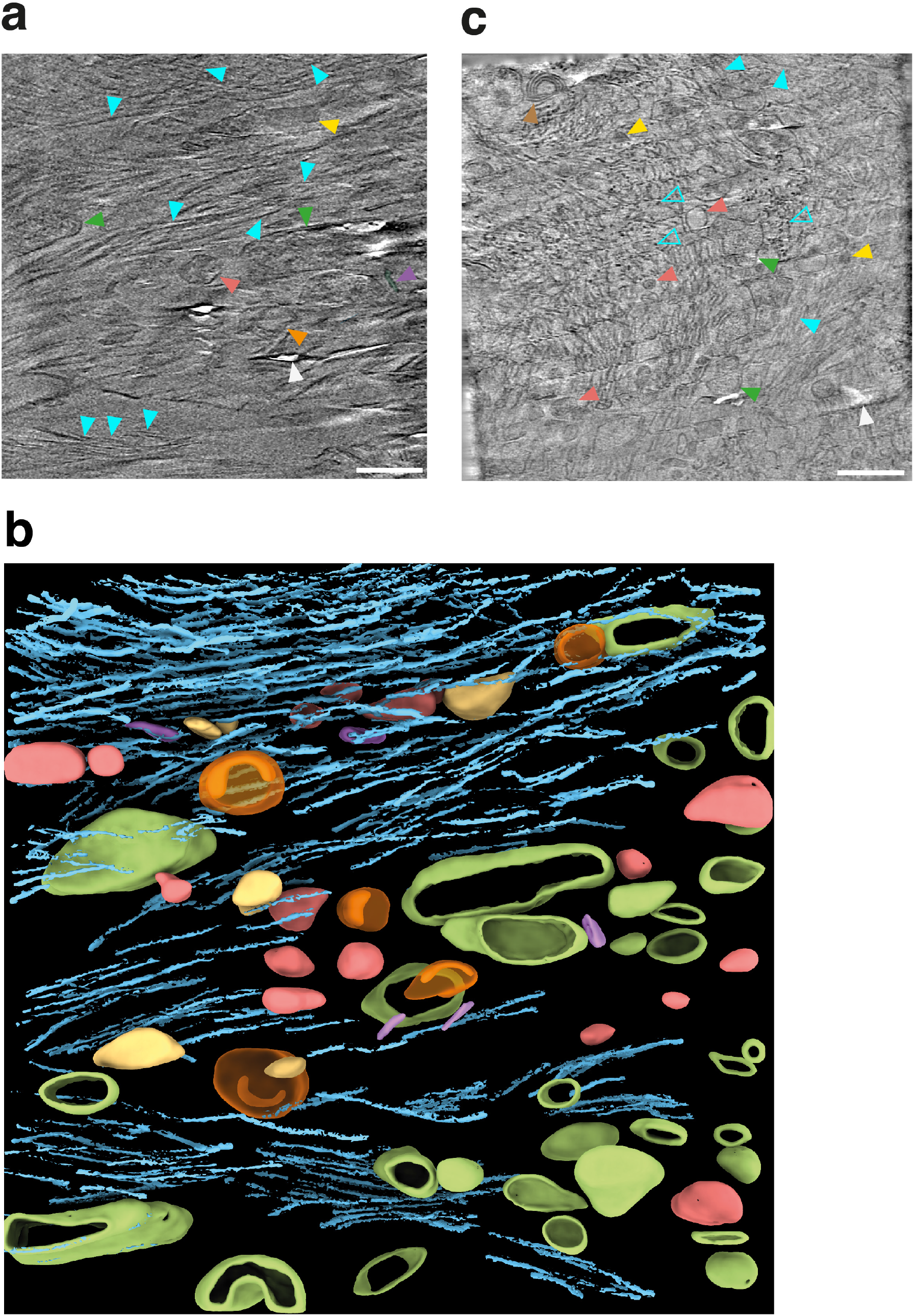
CryoCLEM guided cryo-electron tomography of amyloid plaques showing pathology within central regions of MX04-labelled amyloid plaques. **(a)** Tomographic slice through cryoET reconstruction of amyloid pathology within cortex showing cellular and molecular architecture. Filled and open cyan arrowheads, β-amyloid fibril oriented in the x/y plane and along the z-axis of the reconstructed tomogram, respectively. Dark green arrowhead, cellular compartment. These were larger than the volume of the tomogram and were therefore open at the top or bottom edge of the cryosection. Red arrowhead, spherical exosome. Orange arrowhead, C-shaped exosome. Purple arrowhead, ellipsoidal extracellular vesicle. Yellow arrowhead, extracellular droplet. White arrowhead, localised knife damage in tissue cryo-section. Tomogram was collected with a Volta phase plate. Scale bar, 100 nm. **(b)** 3D segmentation of macromolecular and cellular constituents in a representative tomographic volume of a central region of an MX04-labelled amyloid plaque. Colours as in (**a**). **(c)** Tomographic slice through cryoET reconstruction of pathology collected. Arrowheads as in **a**, except with the addition of filled and open cyan arrowheads, β-amyloid fibril oriented in the along the z-axis of the reconstructed tomogram. Brown arrowhead, multilamellar body. Scale bar, 100 nm. Tomogram was collected without a Volta phase plate. See **Extended Data Fig. 3** showing additional examples of tomographic data collected at central regions of amyloid plaques. See **Supplementary Information Movie 1 and 2** showing example in-tissue tomographic volumes of encompassing central regions of amyloid plaques. See **Methods** for criteria used to identify macromolecular and cellular constituents of tomograms.

**Figure 3.**
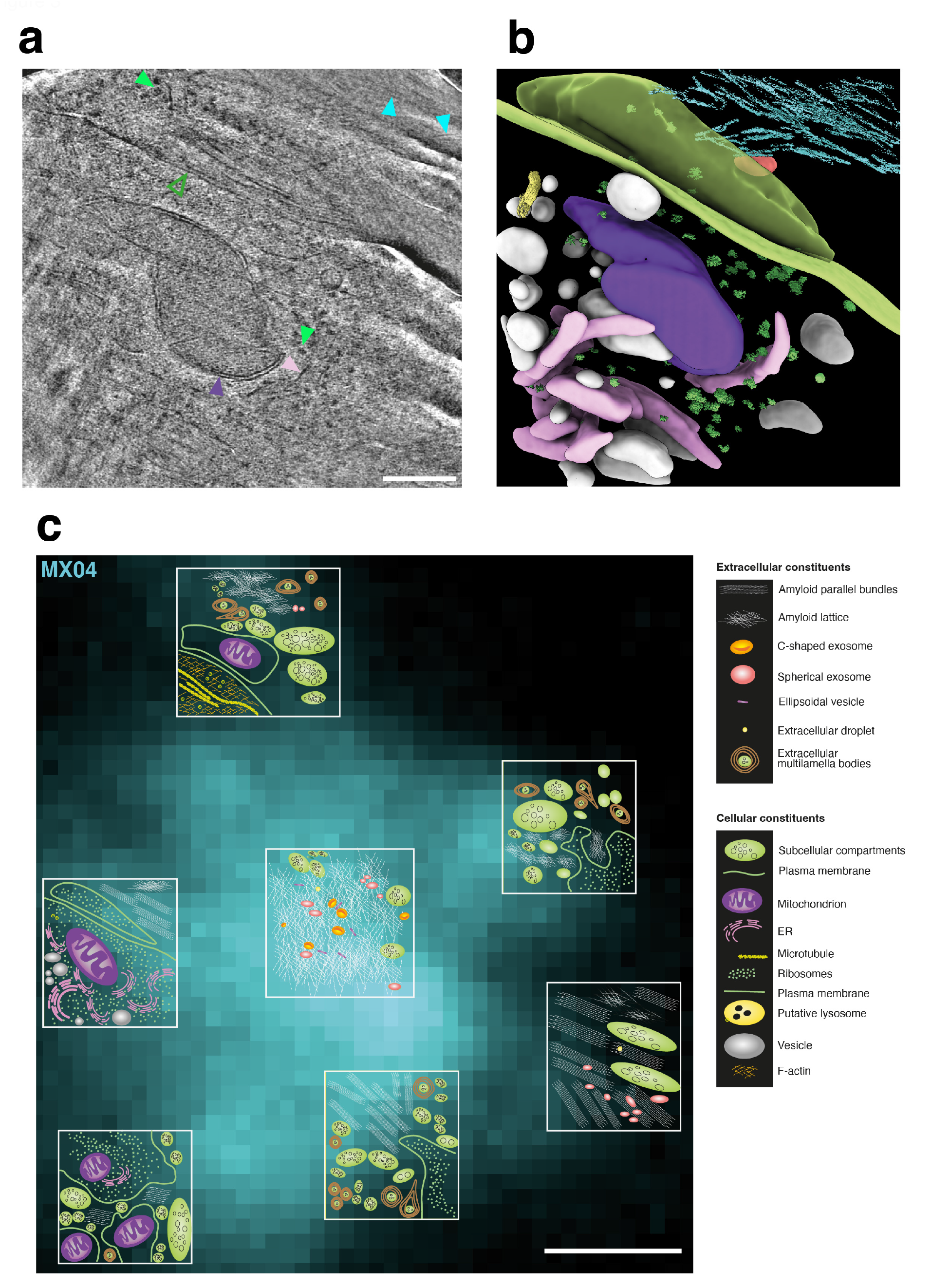
A cryoET survey of the molecular architecture of β-amyloid plaques. **(a)** Tomographic slice through cryoET reconstruction of region at the periphery of MX04-labelled amyloid plaque. Cyan arrowhead, amyloid fibril. Open dark green arrowhead, plasma membrane. Closed light green arrowhead, ribosome. Purple arrowhead, mitochondria. Pink arrowhead, rough endoplasmic reticulum. Scale bar, 100 nm. See also **Extended Data Fig. 4** showing additional examples of tomographic slices of peripheral regions of amyloid plaques. See **Supplementary Information Movie 3 and 4** showing example in-tissue tomographic volumes of encompassing central regions of amyloid plaques. **(b)** 3D segmentation of macromolecular and cellular constituents in a representative tomographic volume of peripheral region of an MX04-labelled amyloid plaque. Extracellular constituents: Cyan, amyloid fibril. Red, exosome. Cellular constituents: Green surface, plasma membrane. Green particles, ribosomes. Purple, mitochondria. Pink, rough endoplasmic reticulum. Yellow, microtubule. White, intracellular membrane vesicles. **(c)** Schematic representation of the native in-tissue 3D molecular architecture of an amyloid plaque overlaid on the MX04 signal of an amyloid plaque. Seven tomograms were collected (yellow boxes) at central and peripheral regions of a single amyloid plaque. The topology of molecular and cellular constituents is shown for each tomogram. See **Extended Data Table 1**.

In central regions of MX04 plaques (**Fig. 2 and Extended Data Fig. 3**) the most abundant non-fibrillar constituents were extracellular vesicles (exosomes). These were readily separable into different types (**Fig. 2, Fig. 3, Extended Data Fig. 3**): i) Spheroidal (50-200 nm diameter) exosomes, ii) exosomes containing a cup or C-shaped membranes in their lumen, and iii) ellipsoidal vesicles (5-20 nm diameter). The prevalence of exosomes within plaques (at least 5 exosomes in 17 of 23 tomograms) is in marked contrast to the parenchyma of healthy tissue where they have rarely been observed^24^. Extracellular droplets were also present in 30% of in-tissue tomograms (**Figs. 2 & 3**). These were smooth, 80-120 nm diameter spheroidal structures, similar to lipid droplets, but are five-to ten-fold smaller than the lipid droplets that reside intracellularly within healthy cells^25^. Extracellular multilamella bodies composed of vesicles wrapped in multiple concentric rings of lipid membrane were also apparent in 30% of tomograms (**Fig. 2c and Fig. 3c**). Similar intracellular intermediates of autophagy have been detected by conventional EM of AD brain^26^.

At the periphery of MX04-stained plaques, subcellular membrane-bound compartments interdigitated or wrapped around amyloid were observed (**Fig. 3a-c and Extended Data Fig. 4**). The cytoplasm of these cells contained ribosomes tethered to rough endoplasmic reticulum, as well as mitochondria and microtubules. These architectures are consistent with the microglial and astrocytic cells that typically surround amyloid plaques^8^. There was no indication of MX04-labelled β-amyloid within the cytoplasm of these cells within or near the amyloid plaques. To verify that *App*^*NL-G-F*^ amyloid is extracellular, we labelled tissue using an extracellular fluorescent marker, dextran-AF647. CryoCLEM of cryo-sections from these samples indicated MX04 overlapping with dextran-AF647, confirming the extracellular location of amyloid in these tomographic volumes (**Extended Data Fig. 5**).

**Figure 4.**
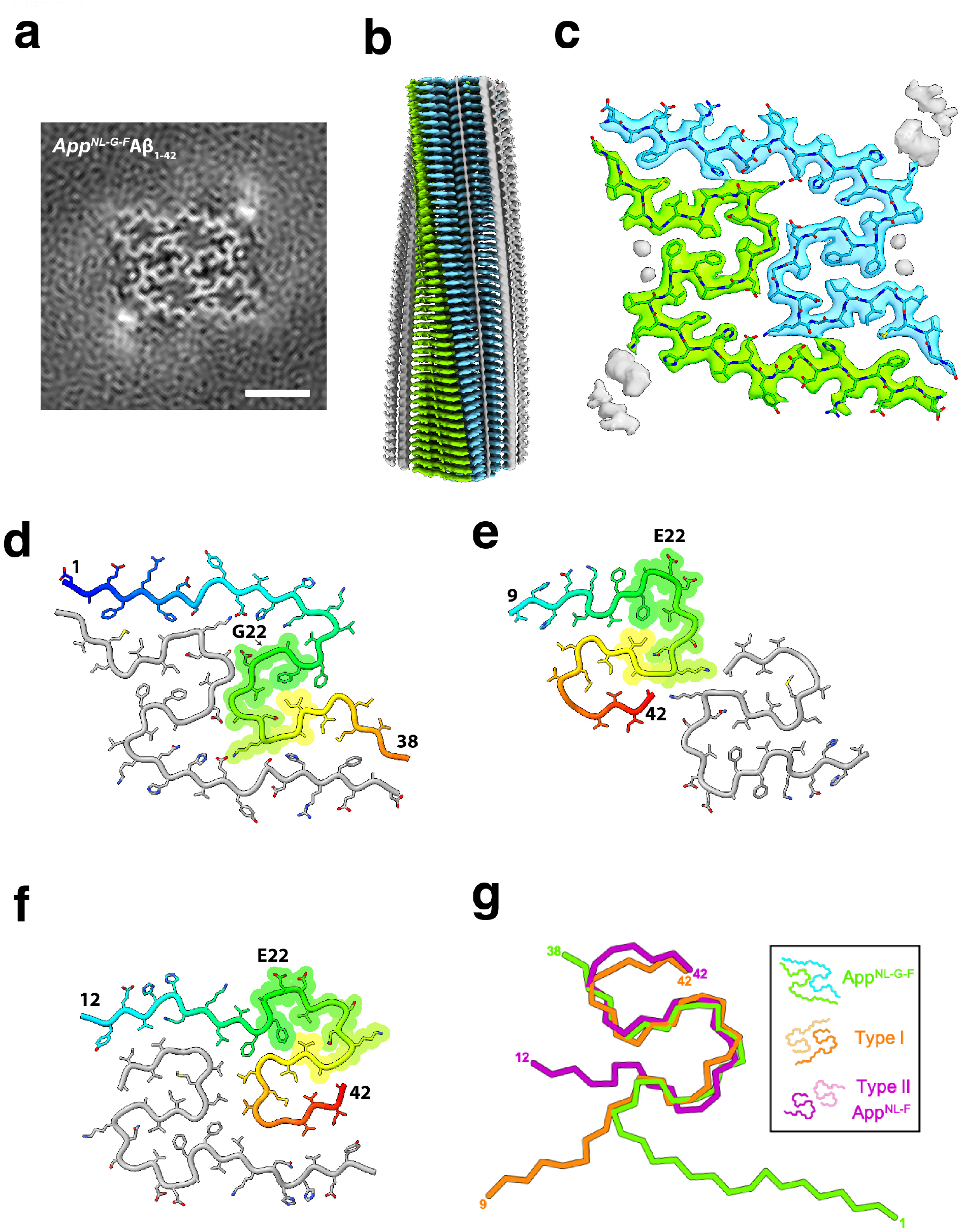
The atomic model of *App*^*NL-G-F*^ Aβ amyloid determined by single-particle cryoEM reconstructions from sarkosyl-extracts. **(a)** Central section through the unmasked final *App*^*NL-G-F*^ amyloid cryoEM map representing a single helical layer (∼4.8 Å, 6x slices averaged) determined at a resolution of 3.0 Å (FSC 0.143). Scale bar, 2.5 nm. **(b)** Perpendicular view of the cryoEM map along the fibril axis, colour-coded by the protein chain (cyan and green), with unmodelled associated density in grey. The image shows clear resolution of individual β-strand layers within the amyloid core. **(c)** The cryoEM density and fitted model sectioned around a single helical layer, colour-coded as in (**b**), with unmodelled associated density in grey. **(d)** One layer of the *App*^*NL-G-F*^ β-amyloid model with one subunit coloured blue-to-red from N-to-C-terminus (excluding disordered residues 39-42). **(e)** Type II Aβ_42_ fibril model from human AD brain (PDB: 7Q4M)^13^, with each subunit coloured blue to red (N-to C-terminus). Note that residues 1-38 are ordered in the Arctic structure, while residues 9-42 are structured in the type II fibrils. **(f)** Type I Aβ_42_ fibril model (PDB: 7Q4B)^13^, coloured as in **e**. Note that residues 1-38 are ordered in the Arctic Aβ structure while residues 12-42 are structured in type I fibrils. **(g)** Superposition of the subunit folds from each of the three displayed fibril models reveals a common protein conformation between residues 19-36, with diverging terminal folds. The per atom RMSD values for residues 19-37 between the models is: 1.59 Å for *App*^*NL-G-F*^ vs Type I and 1.36 Å for *App*^*NL-G-F*^ vs Type II fibrils.

**Figure 5.**
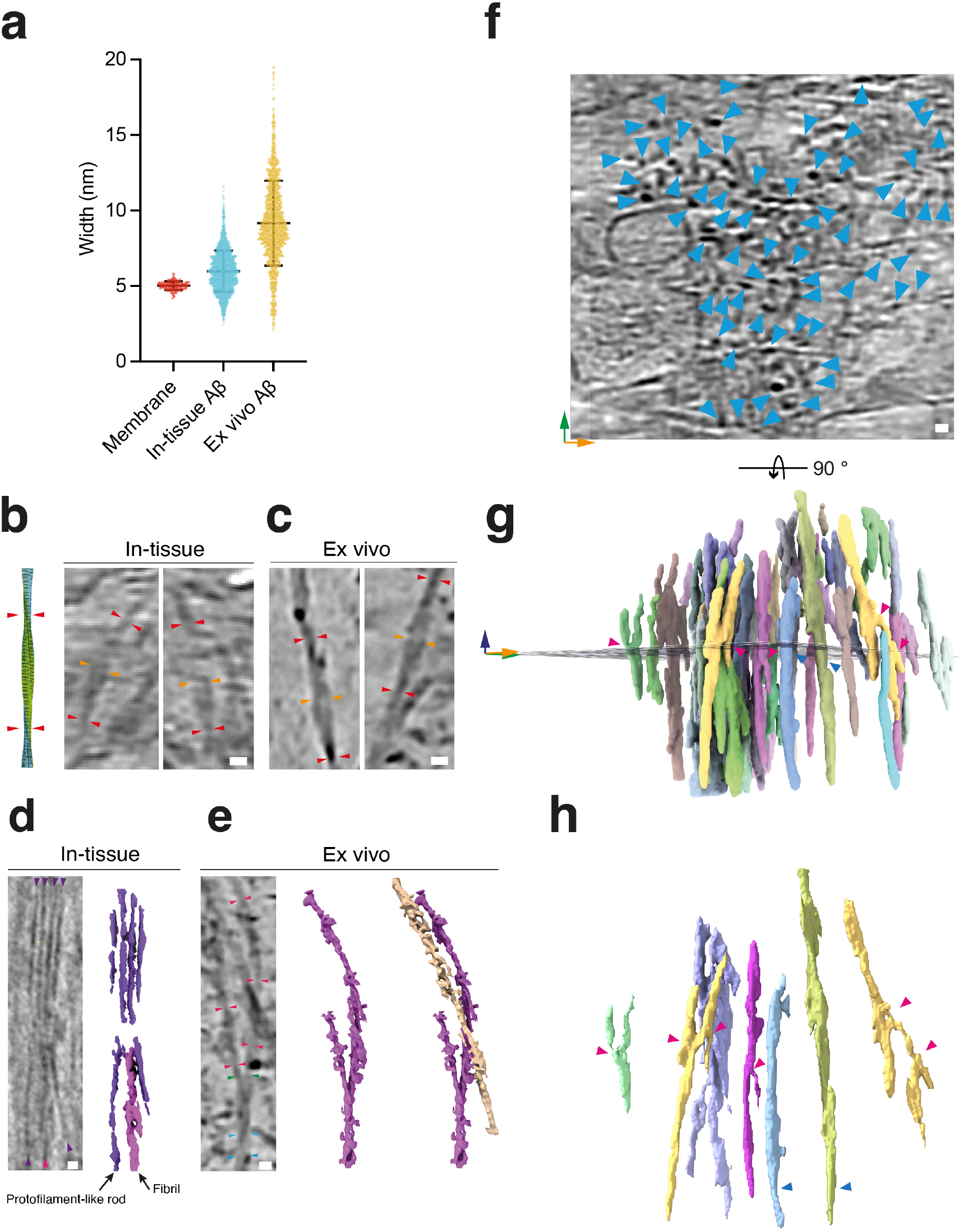
A diversity of fibril types within amyloid plaques. **(a)** Scatterplot showing the distribution of lipid membrane (n=541) and fibril width from in-tissue (cryoET volumes of cryo-sections from *App*^*NL-G-F*^ amyloid plaques, n=1840) and ex vivo (cryoET of sarkosyl-extracted amyloid fibrils, n=1470). Middle and top/bottom clack bars indicate mean and one standard deviation. **(b)** Left, atomic model Arctic Aβ fibril. *Right*, tomographic slices showing *in-tissue* with an apparent helical twist of two protofilaments. Red arrowheads, apparent crossover. Orange arrowheads, point of maximum fibril width. Scale bar, 10 nm. **(c)** Same as **b** but for ex vivo amyloid. Scale bar, 10 nm. (d) *Left*, tomographic slice, and *right*, tomographic density showing in-tissue amyloid fibrils with 3-4 nm width fibrils (purple arrowheads) and a 7-10 nm fibril (magenta arrowhead). Scale bar, 10 nm. **(e)** *Left*, tomographic slice of branched fibrils in *ex vivo* amyloid. Blue arrowheads, 7-10 nm fibril. Green arrowheads, fibril branch point. Magenta arrowheads, protofilament. Scale bar, 10 nm *Middle*, tomographic density of branched fibril. *Right*, Same as *Middle* except with nearby 7-13 nm fibril (gold coloured) included. See **Extended Data Fig. 9c-f** showing additional examples of branched fibrils in *ex vivo* amyloid. See also **Supplementary Information Movie 5** showing tomographic volume of ex vivo fibrils. **(f)** Tomographic slice of in-tissue amyloid with fibrils oriented on the z-axis of reconstructed tomographic volume. Cyan arrowhead, high-contrast spot corresponding to a single fibril. Orange and green arrows, *x*- and *y*-axis of tomogram, respectively. Scale bar, 10 nm. **(g)** Segmented tomographic density of fibrils shown in **c**, viewed from the side. Fibrils are coloured by connectivity. Magenta arrowhead, putative fibril branch points. Blue arrowhead, unbranched fibril. Orange, green, and blue arrows, *x*-, *y*-, *z*-axis, respectively. **(h)** Raw tomographic density of a subset of fibrils from panel (**g**). Magenta arrowhead, putative branch points. Blue arrowhead, unbranched fibril. See **Extended Data Fig. 10** showing an additional example of in-tissue branched amyloid. See **Supplementary Information Movie 2** showing parallel bundles of fibrils oriented on the z-axis of the in-tissue tomographic volume.

### The high resolution structure of Arctic Aβ fibrils

Recent structures of Aβ amyloid fibrils extracted from post mortem human AD brain revealed two different fibril forms (type I and II)^13^. The same study showed that fibrils from *App*^*NL-F*^ mice that express wild-type Aβ_1-42_ form type II Aβ fibrils^13^. The structure of amyloid fibrils that result from the *App*^*NL-G-F*^ knockin strain, which contains the Arctic mutation (E22G) within the Aβ_1-42_ peptide, have not been reported. We therefore performed a sarkosyl-extraction of Aβ fibrils from *App*^*NL-G-F*^ mouse forebrains, for single-particle cryoEM, using procedures developed by Yang and co-workers^13^. The sample contained fibrils with an overt helical twist corresponding to a predominant crossover distance of ∼66 nm (**Extended Data Fig. 6a**), allowing us to determine a 3.0 Å resolution structure of the fibrils from *App*^*NL-G-F*^ mouse brain (**Fig. 4a, b and Extended Data Fig. 6**). The fibril is two-fold symmetric about the helical axis, allowing us to unambiguously build in two identical copies of residues 1-38 of the Aβ_1-42_ sequence into the cryoEM density map (**Fig. 4c**). We also observed a minor population of wider fibril segments composed of an apparent dimeric assembly of fibrils containing four copies of Aβ per molecular layer that had too few particles to obtain a helical solution **(Extended Data Fig. 6c,d)**. Consistent with earlier MS imaging^27^, mass spectrometry of *ex vivo* purified amyloid identified mainly Aβ_1-42_ and smaller peaks corresponding to Aβ_1-38/39_ Aβ_11-42_ and Aβ_1-40_ (**Extended Data Fig. 7a,b**), suggesting that ∼61% C-terminal residues 39-42 of Aβ_1-42_ are present in the fibrils, but are disordered in our map. The *App*^*NL-G-F*^ *ex vivo* fibrils displayed a different Aβ fold than that found in fibrils extracted from *App*^*NL-F*^ mice and AD brain, which lack the Arctic E22G mutation. The backbone conformation of the Aβ peptides are arranged as two ‘S’-shaped protofilaments that form an extensive inter-protofilament interface. The ordered core of each protofilament contains residues 1-38, compared to residues 9-42 in type I and II wild-type Aβ_1-42_ ^13^. The solvent-accessible surface is also fundamentally different (**Extended Data Fig. 7c**). In both type I and II fibrils formed from wild-type Aβ_1-42_, E22 is surface exposed^13^, whereas the equivalent position in Arctic Aβ, residue G22, is buried at the protofilament interface (**Fig. 4d-f**). Thus, it appears likely that the wild-type Aβ would be unable to adopt the Arctic Aβ amyloid fold because of steric clashes. Alignment of available Aβ_1-42_ structures from post mortem human AD brains, and the two distinct Aβ amyloid structures from *App*^*NL-F*^ and *App*^*NL-G-F*^ mice, indicated that residues 19-37 form the same structure (1.35 Å and 1.95 Å Cα RMSD A*pp*^*NL-G-F*^ with Type I and Type II fibrils, respectively, **Fig. 4g**). This conserved structural element adopts different positions with respect to the fibril axis and the interface between symmetry related protofilaments, giving rise to the different amyloid structures observed. The N- and C-terminal residues outside of this structural element adopt different conformations, wrapping around this structural element, wrapping around that of its neighbouring, symmetry-related protofilament, or remaining unstructured, in these different amyloid folds.

### Protofilaments and branched fibrils in amyloid plaques

How well the structure of sarkosyl-extracted fibrils represents the array of fibrils observed *in situ* using cryoET was next addressed by collecting 27 tomograms of the sarkosyl extracted fibrils (**Extended Data Fig. 7d**) and directly comparing the widths of fibrils in both *ex vivo* and in-tissue cryoET datasets (**Fig. 5a**). To rule out that MX04 altered the structure of the in-tissue fibrils, we also determined the structure of fibrils purified from MX04-injected *App*^*NL-G-F*^ mice using single particle cryoEM. The results revealed a structure that was indistinguishable from that obtained using unlabelled β-amyloid (**Extended Data Fig. 8a**). The width of lipid membrane bilayers served as an internal ‘molecular ruler’ of the accuracy of distance measurements within in-tissue tomograms, yielding an average thickness of 5.01 ± 0.38 nm (mean width ± SD; **Fig. 5a**), as expected^28^. The average width of *ex vivo* purified fibrils measured in tomographic slices (9.2 ± 2.8 nm mean ± SD fibril width) (**Fig. 5a**) was consistent with the width of the reprojected 3D atomic structure of the fibrils determined using cryoEM (8.5 ± 0.95 nm mean ± SD fibril width; **Extended Data Fig. 8b**). Unexpectedly, the average width of in-tissue fibrils was significantly smaller than that of *ex vivo* fibrils in tomographic volumes (∼6.0 ± 1.4 nm versus 9.2 ± 2.8 nm mean ± SD fibril width, respectively; P<0.0001). This difference could indicate a variation in the proportions of distinct fibril populations.

To assess further the possibility of distinct fibril populations we tested whether fibrils of different width were evenly distributed across amyloid plaques by performing one-way analysis of variance, which indicated that there was a significant enrichment of thin versus thick fibrils in different in-tissue tomograms (**Extended Data Fig. 9a**; One-way ANOVA: F(df=11)=39.13, P=2×10^−16^). This suggested the variance of fibril width is region-specific. Importantly, all in-tissue tomograms showed an MX04 cryoCLEM signal in the location of regions enriched in either thin or thick fibrils, suggesting that both are composed of amyloid. However, fibril widths did not segregate on the basis of their location at the periphery or in the core of amyloid plaques (**Extended Data Fig. 9b**).

Closer examination of the population of thick (7-11 nm width) fibrils in less crowded regions of in-tissue and *ex vivo* tomograms showed apparent crossovers (**Fig. 5b, c**). The helical pitch of these fibrils, corresponding to the distance between crossovers of *ex vivo* and in-tissue fibrils, was similar (65.1 ± 2.4 nm mean ± SD versus 65.9 ± 1.6 mean ± SD helical pitch, respectively) and consistent with that observed in the single particle reconstruction (**Fig. 5b**; 65.5 nm helical pitch). In contrast, the thin (3-5 nm width) fibrils lacked the characteristic crossover of *ex vivo* Aβ atomic structures (**Fig. 5d**). Since the width of 3-5 nm fibrils was half that of the Aβ fibril structure, we describe this form of amyloid as ‘protofilament-like rods’. Closer inspection of this minority of protofilament-like rods in *ex vivo* tomograms revealed that they were found branching away from a fibril (**Fig. 5e and Extended Data Fig. 9c-f**). Some branch points gave rise to protofilament-like rod extensions that were thinner (3-5 nm width) than their parent fibril (7-11 nm width), whereas other branches connected two 7-11 nm fibrils (**Extended Data Fig. 9c)**.

To explore whether amyloid branching exists within in-tissue tomograms, we examined the raw tomographic density of parallel fibril bundles oriented along the tomographic *z*-axis. These provided sufficient contrast to trace individual fibrils throughout the tomographic volume (**Fig. 5f**). These segmented tomographic volumes showed branched fibrils intermingled with unbranched fibrils within amyloid plaques (**Fig. 5g, h and Extended Data Fig. 10**). Thus, in-tissue tomographic data suggest that in addition to Aβ fibrils containing two, intertwined protofilaments with the Aβ Arctic amyloid fold, amyloid plaques also contain protofilament-like rods and branched amyloid structures.

## Discussion

Tomograms of Aβ amyloid in-tissue describe a complex molecular architecture of β-amyloid plaques that includes fibrils, protofilament-like rods and branched amyloid. This diversity of amyloid architecture occurs alongside non-amyloid constituents, including exosomes, extracellular multilamellar bodies and cellular compartments, consistent with earlier observations using conventional EM of resin embedded post mortem AD tissues^7,8^, animal models^9^, and tomography of resin-embedded *in vitro* cell-based model of Aβ fibril toxicity^23^. The detail afforded by cryo-preservation of native tissue and cryoET suggests the existence of distinct types of exosomes, including exosomes containing a luminal C-shaped membrane. Extracellular droplets that resemble lipid droplets were also observed. These had eluded identification in earlier studies^7–9,23^. All of the non-amyloid extracellular constituents are rarely observed in normal brain tissue^22,24^, suggesting that these features are indicative of pathology, perhaps related to earlier stages of amyloid plaque biogenesis^23,29^, or an on-going cellular response to the accumulation of amyloid^30^.

Using cryoEM we determined the native in-tissue 3D molecular architecture of extracellular amyloid plaques of the Arctic variant of Aβ in the mammalian brain and atomic models of *ex vivo* amyloid purified from these tissues. These data show that the fibrils of the Arctic variant have a unique fold, that differs from those observed previously for fibrils purified for wild-type Aβ_1-40_ from CAA brain^15^, wild-type Aβ_1-42_ fibrils Type I and Type II purified from human AD brain post mortem and FAD mouse models as well as those generated *in vitro* from purified Aβ_1-42_ peptide^12,13^. The structural differences include the identity of the residues involved in the amyloid core and in the arrangement of the two protofilaments, which nonetheless, each contain a common structural element involving residues ∼19-36 of the 42 residue sequence. These structural differences, exposing different residues on the surface of Arctic Aβ fibrils (**Extended Data Fig. 7c**), could explain the inability of the diagnostic reagent Pittsburgh B to detect Arctic amyloid in PET imaging of patients^31^ and animal models^32^.

The single-particle reconstruction of Arctic Aβ fibrils describes the structure of the major species of amyloid in *App*^*NL-G-F*^ mice. However, cryoET data collected from *ex vivo*, purified amyloid also revealed the existence of rare protofilaments and branched fibrils. The presence of protofilament-like rods and fibril branch points was also apparent within in-tissue tomograms of *App*^*NL-G-F*^ amyloid plaques. Protofilaments and branched fibrils have also been observed previously by atomic force microscopy of *in vitro* Aβ_1-42_ assembly assays^33^, albeit not all reported *in vitro* preparations of Aβ have shown this architecture^34^. Short protofilaments branching from fibrils have been observed by cryoEM imaging of *in vitro* Aβ aggregation assays^35^. The existence of protofilament-like rods and branched fibrils *in situ* provides a structural model for the dense architecture of plaque cores. Such branch points could result from catalysis of fibril growth *in vivo* by secondary nucleation^35,36^ and provide a structural model for the focal accumulation of Aβ within amyloid plaques.

The structural diversity of Aβ is recognised as one of the challenges in developing therapeutics to treat Alzheimer’s disease. Importantly, antibodies developed as an immunotherapy and tested in recent clinical trials were raised against the Arctic Aβ variant, which on the basis of *in vitro* prepared Aβ assemblies was expected to generate a high concentration of small, worm-like protofibrils^17,37^. Our in-tissue tomographic data of Arctic Aβ from murine FAD models reveal no such enrichment of short protofibrils. However, it is possible that other *in situ* structures of Aβ, including protofilament-like rods and branched amyloid contain equivalent epitopes to *in vitro* prepared protofibrils that mediate the effect of Arctic Aβ-directed immunotherapies. Future studies are needed to explore further the origin and detailed structural differences of these amyloids and to understand the in-tissue molecular architectures of post mortem AD brain.

## Acknowledgements

We are grateful to Takaomi Saido (RIKEN Centre for Brain Sciences) for generously providing us with the *App*^*NL-G-F*^ mouse line. We would like to thank Rebecca Thompson, Emma Hesketh, Martin Fuller, Daniel Maskell, and Yehuda Halfon for help maintaining and setting up the Astbury Biostructure Laboratory (ABSL) cryoEM facility, including cryo-preservation/imaging and Titan Krios microscopes. We are grateful to Andrew Horner, Ilona Rigo and Melanie Reay for technical support and Alex Taylor and Liam Aubrey for enlightening discussions.

## Author Contributions

C.L., A.B. and R.A.W.F. organised breeding, analysed tissue, established the workflow and collected data for in-tissue cryoCLEM/cryoET. C.L., A.B., S.D. and R.A.W.F. reconstructed and analysed in-tissue cryoET data. S.G., M.W. and R.A.W.F. prepared and analysed *ex vivo* amyloid. M.W. collected and performed cryoEM structure determination. Y.X. performed and analyzed mass spectrometry of *ex vivo* amyloid. M.W., and C.L. collected reconstructed *ex vivo* amyloid cryoET data. M.W., N.A.R., and R.A.W.F. analysed *ex vivo* amyloid cryoET. R.A.W.F., N.A.R., and S.E.R. supervised the project. All authors contributed to writing the manuscript.

## Supplementary Information Figure legends

**Movie 1**. Video showing tomographic volume at central region of MX04-labelled amyloid plaque. Related to **Fig. 2a**.

**Movie 2**. Video showing tomographic volume at central region of MX04-labelled amyloid plaque. Related to **Fig. 2b**.

**Movie 3**. Video showing tomographic volume at the peripheral region of MX04-labelled amyloid plaque. Related to **Fig**. 3a.

**Movie 4**. Video showing tomographic volume at the peripheral region of MX04-labelled amyloid plaque. Related to **Fig. 3a**.

**Movie 5**. Video showing tomographic volume of ex vivo purified amyloid. **Related to Extended Data Fig. 5e**.

## Methods

### Laboratory Animals

Two 11-12 month-old *App*^*NL-G-F*^ knockin mice^16^ were used for in-tissue structural analysis by cryoCLEM and cryoET. 11-13 month-old *App*^*NL-G-F*^ and wildtype controls we used for immunohistochemistry. Two 11-13 month-old *App*^*NL-G-F*^ knockin mice were used for each *ex vivo* amyloid purification. Laboratory animals were housed and bred according to British Home Office Regulations, local ethical approval, and NIH guidelines.

### Preparation of acute brain slices

Mice received an i.p. injection of 5 mg/kg Methoxy-X04 (Tocris) in 10% w/v DMSO containing phosphate-buffered saline. 24 hours after injection, mice were injected with pentobarbital and intracardially perfused with N-methyl-D-glucamine (NMDG)-HEPES solution (93 mM NMDG, 2.5 mM potassium Chloride, 1.2 mM sodium hydrogen carbonate, 20 mM HEPES, 25 mM glucose, 5 mM sodium ascorbate, 2 mM thiourea, 3 mM sodium pyruvate, 10 mM magnesium sulphate heptahydrate, 0.5 mM calcium chloride dihydrate, osmolality = 300 - 315 mOsmol/kg, pH 7.4)^38^. Brains were retrieved and 100 μm thick acute slices were prepared using a vibratome (speed 0.26 mm/s, Leica, VT1200S, Campden Instruments Limited blades, J52/11SS blades) in ice-cold carboxygenated NMDG-HEPES solution.

### Immunohistochemistry and confocal fluorescence microscopy

Free-floating acute brain slices were fixed in 4% paraformaldehyde (PFA), blocked in 5% BSA, 0.1% w/v Triton-X100 containing Tris Buffered Saline (TBS, 50 mM Tris-Cl, 150 mM NaCl) and after three washes with TBS, incubated with 6E10 anti-amyloid-beta 1-16 mouse IgG1 (1:750, Biolegend, 803001) in 0.1% w/v Triton-X100 containing TBS at 4°C for 24 h. After three washes in TBS, slices were incubated with anti-mouse-IgG1-AF-633 (1:1000, Lifetechnologies, A21126) in 0.1% w/v Triton-X100 in TBS for 48 h at 4°C. After 3 washes, slices were mounted in Vectashield (Vector Laboratories) on superfrost slides (Erpredia, J1810AMNZ) with coverslips (Academy, 0400-8-18). Images were captured with a confocal laser scanning microscope (Zeiss LSM 700) using a 10/0.3 and a 20x/0.5 numerical aperture (NA) air objective lens, with frame size 1024×1024 pixels and 512×512 pixels, respectively (Methoxy-X04 excitation: 405nm, emission; 435 nm; AF-633 excitation: 639nm, emission; 669 nm).

### High pressure freezing

Acute brain slices were sampled by collecting 2 mm diameter cortical tissue biopsies. These were incubated in cryoprotectant (20% w/v 40,000 Dextran^22^ in NMDG-HEPES solution) for ∼45 min at RT. 100 μm deep wells of the specimen carrier type A (Leica, 16770152) were filled with cryoprotectant, tissue biopsies were carefully placed inside and covered with the flat side of the lipid-coated specimen carrier type B (Leica, 16770153) and high-pressure frozen (∼2000 bar, - 188°C) using a Leica EM ICE.

### Cryo-ultramicrotomy

High pressure frozen sample carriers were imaged with a cryo-fluorescence microscope (cryo-FM, Leica EM Thunder with HC PL APO 50/0.9 NA cryo-objective) at −180°C to determine the location of Aβ plaques. The brightest fluorescent signal with an excitation of 350/50 nm and emission of 460/50 nm underneath the surface of ice were chosen for cryo-sectioning. The distance from Aβ plaque of interest to the edge of the carrier was measured to target the collection of cryosections containing an amyloid plaque. Next, the carriers were transferred to a cryo-ultramicrotome (Leica EM FC7, −150°C) equipped with trimming (Trim 45, T1865 and trim 20, T399) and CEMOVIS (Diatome, cryo immuno, MT12859) diamond knives. A 100 × 100 × 60 μm trapezoid stub of tissue was trimmed that contained the target amyloid plaque, from which 70 - 150 nm thin sections were cut at −150°C with a diamond knife (Diatome, cryo immuno, MT12859). Sections were picked up with a gold eyelash and adhered onto a glow discharged (Cressington glow discharger, 60 s, 10-4 mbar, 15 mA) 1.2/1.3 or 3.5/1, 300 mesh Cu grid (Quantifoil Micro Tools) using a Crion electrostatic gun^39^ and micromanipulators^40^.

### Cryogenic fluorescence microscopy

High pressure frozen tissue inside gold carriers and tissue cryo-sections were screened for fluorescence using a cryogenic fluorescence microscope Leica EM Thunder with a HC PL APO 50x/0.9 NA cryo-objective, Orca Flash 4.0 V2 sCMOS camera (Hamamatsu Photonics) and a Solar Light Engine (Lumencor). A DAPI filter set (excitation 365/50, dichroic 400, emission 460/50) was used to detect methoxy-X04 labelled amyloid. A rhodamine filter set (excitation 546/10, dichroic 560, emission 525/50), was used as a control imaging channel. The images were acquired with a frame size of 2048×2048 pixels. Tile scans of high-pressure frozen carriers were acquired with 17% laser intensity for 0.1 s. Z-stacks of ultrathin cryo-sections were acquired with 30% intensity and an exposure time of 0.2 s. Images were processed using Fiji ImageJ^41^.

### Cryogenic correlated light and electron microscopy (cryo-CLEM)

The location of amyloid plaques in ultrathin cryo-sections was assessed by cryogenic fluorescence microscopy based on Methoxy-X04 fluorescence (excitation 370 nm, emission 460-500 nm). Grid squares that contained a signal for Methoxy-X04 were selected for electron tomography. The alignment between cryoFM images and electron micrographs was performed using a Matlab script^42,43^, in which the centres of 10 holes in the carbon foil surrounding the region of interest were used as fiducial markers to align the cryo-FM and cryo-EM images.

### Cryo-electron tomography imaging and reconstruction

Electron microscopy was performed with a ThermoFisher 300 keV Titan Krios G2, X-FEG equipped with Gatan K2 XP summit direct electron detector and BioQuantum energy filter, in the Astbury Biostructure Laboratory (ABSL) at the University of Leeds. Tomographic tilt series were collected from +60° to −60° in 2° increments using a dose symmetric tilt scheme^44^ in serialEM^45^. Two of the datasets were collected with a Volta phase plate^46^ conditioned to 0.4 - 0.7 Π rad with a 0.8 to 1.3 μm defocus. A further two datasets were collected with 70 μm objective aperture and 5.0 to 6.5 μm defocus. Each tilt increment received 2 s exposure (fractionated into 8 movie frames) at a 0.5 e-.Å^2^.s^-1^ dose rate, resulting in a total dose per tilt series of ∼61 electrons and pixel size of 3.42 Å. Dose fractions were aligned and tomograms were reconstructed using patch tracking in IMOD^47^. Segmentation was performed using the coordinates of fibrils manually picked in IMOD and of the lipid bilayer of membranes in Dynamo^48^. Figures were prepared in ChimeraX^49,50^ using SIRT reconstructed tomograms that were deconvoluted with Isonet^51^ using defocus values estimated with Gctf^52^.

### Annotation and analysis of macromolecular constituents in tomograms

Annotation described in Extended Data Table 1 was performed blind by two curators to catalogue the constituents of in-tissue tomograms. First, each curator annotated all SIRT reconstructed tomograms independently. Next, a third curator inspected and certified each annotation (Extended Data Table 1). The following constituents were identified: 1) Amyloid fibrils were assigned on the basis of methoxy-X04 cryoCLEM labelling and rod-shape within extracellular locations of the tissue. 2) Extracellular vesicles (exosomes) were defined as membranes that were closed and situated with extracellular locations. Exosomes were subdivided into three structural categories. 2a) Spherical exosomes (50-200 nm diameter) enclosed by a single plasma membrane. 2b) C-shaped exosomes composed of cup-shaped membrane within the lumen of a spherical exosomes. 2c) Ellipsoidal vesicles (5-20 nm diameter). 4) Extracellular droplets: 80-120 nm amorphous and smooth spheroidal particles that resembled lipid droplets. 7) Extracellular multilamella bodies: contained 60-200 nm vesicle or subcellular compartment rapped in a spiral of membrane lipid bilayer. 5) Subcellular compartments: containing a higher tomographic density than the extracellular space that is characteristic of the high concentration of proteins in the cell cytoplasm compared to the extracellular space. 6) Cell plasma membranes: marked the boundary between the higher tomographic density of the cytoplasm and the lower tomographic density of extracellular interstitial space. 7) Mitochondria: defined by the double membrane including outer membrane and inner mitochondrial cristae. 8) Ribosomes were distinguishable as ∼30 nm particles with higher tomographic density than surrounding macromolecules, in accordance with their high nucleic acid content. 9) Rough endoplasmic reticulum: intracellular tubular membranes with ribosomes associated on the outside edge. 10) Microtubules: were identified as ∼25 nm wide filaments. 11) Knife damage: Tissue cryo-sections contained small regions in which the sample has been compressed, leaving a crevasse in the tissue that were readily identified as holes within the tissue^53^.

### Ex vivo purification and single-particle cryoEM data collection of amyloid from App^NL-G-F^ mouse

Two 11-14 month-old homozygous *App*^*NL-G-F*^ knockin mouse forebrains (cortex and hippocampus) were used for each preparation of *ex vivo* amyloid following the sarkosyl extraction protocol reported by Yang and co-workers^12^. The sarkosyl-extracted *App*^*NL-G-F*^ amyloid, both with and without methoxy-X04 pre-treatment, were diluted one in three with EM buffer (20 mM Tris pH 7.4, 50 mM NaCl) and 4 μl applied to 60 s plasma cleaned (Tergeo, Pie Scientific) Quantifoil R1.2/1.3 grids were plasma cleaned (Tergeo, Pie Scientific). Grids were blotted and plunge-frozen in liquid ethane using a Vitrobot Mark IV (FEI) with a 1 s wait and 5 s blot time respectively. The Vitrobot chamber was maintained at close to 100% humidity and 6°C. The single-particle datasets were collected at ABSL (Leeds) using a Titan Krios electron microscope (ThermoFisher) operated at 300 keV with a Falcon4 detector in counting mode (Nominal magnification of 96,000x and 0.83 Å/pixel). A total of 2,428 movies were collected with a nominal defocus range of −1.6 to −3.1 μm and a total dose of ∼52 e-/Å^2^ over an exposure of 8 s, corresponded to a dose rate of ∼4.5 e-/pixel/s. The control methoxy-X04 dataset was collected with an additional Selectris energy filter with a 10 e^-^V slit, a nominal magnification of 130,000x and a pixel size of 0.94 Å. A total of 4,165 movies were collected with a nominal defocus range of −1.4 to −2.9 μm and a total dose of ∼41 e-/Å^2^ over an exposure of 6 s, corresponded to a dose rate of ∼6.1 e-/pixel/s.

### Single particle helical reconstruction of App^NL-G-F^ ex vivo Aβ fibrils

Raw EER movies were compressed and converted to TIFF using RELION-4^54^, regrouping frames into 40 fractions to give a dose per frame of 1.3 e^-^/Å^2^ for the *App*^*NL-G-F*^ dataset and 1.0 e^-^/Å^2^ for the methoxy-X04-treated dataset, respectively. The TIFF stacks were aligned and summed using motion correction in RELION-4 and CTF parameters were estimated for each micrograph using CTFFIND4^55^. Fibrils from roughly 100 micrographs were picked manually and used to train a separate picking model for each dataset in crYOLO^56^ for automated picking with an inter-box spacing of 3x helical repeats (∼14 Å). Micrographs not containing fibrils were removed by screening the lowpass-filtered micrographs generated by crYOLO. This left 380 fibril-containing micrographs (**Extended Data Fig. 6a**) for further processing, from which 63,680 fibril segments were extracted 3x binned with 750 Å box dimensions. Two rounds of 2D classification were performed to remove picking artefacts, generating a cleaned dataset of 46,215 segments of fibrillar classes. The resulting class averages were split into two subsets based on fibril morphology, with 97% selected in the major subset and 3% showing wider fibrils in the minor subset (**Extended Data Fig. 6b** and **6c**). Each subset was re-extracted 2x binned (500 Å box dimensions) and initial models were generated from a respective 2D class average with a helical twist estimate obtained from measured crossover distances using the relion_helix_inimodel2d command^57^. These templates were used to start a one-class 3D classification (initial lowpass filter of 15 Å) to obtain a refined initial model for each form (**Extended Data Fig. 6b** and **6c**). A final 2D classification run on each 2x binned subset was used to remove unfeatured fibril segments. The structure of the wider, minor fibril form could not be accurately determined due to the low starting number of particles (**Extended Data Fig. 6d**). The cleaned 30,070 segments for the major subset were re-extracted unbinned (250 Å box dimensions) for further 3D classification runs with searches of the helical twist (**Extended Data Fig. 6e**). Only one ordered fibril form with a pseudo-2_1_ helical symmetry was identified in the data from several classification attempts. After multiple rounds with increasing angular sampling and initial resolution of lowpass-filter, 2,568 ordered segments were selected and the helical parameters optimised to give good layer separation (**Extended Data Fig. 6f**). The segments refined to 3.3 Å (gold-standard, 0.143 FSC) and then to a final resolution of 3.0 Å after CTF refinement and Bayesian polishing (**Extended Data Fig. 6g** and **6h**). The final map was sharpened using a B-factor of −23 Å^2^ and helical symmetry was applied using the refined helical twist of 179.352° and rise of 2.418 Å respectively. The control, methoxy-X04 treated fibril dataset was similarly processed and yielded an identical structure at a resolution of 3.2 Å, deposited with a sharpening B-factor of −25 Å^2^ (**Extended Data Fig. 7a**). Full details are shown in **Extended Data Table 2**.

Initially, we tried to dock the human Aß42 AD type I fibril structure (PDB: 7Q4B) into the map, which has a superficially similar structure. However neither the overall model nor the register of the amino acid side chains fitted. We therefore built a *de novo* model for one chain using Coot^58^ using the large bulky side chain densities, as well as the regions without side chain density (glycine residues) and general chemistry of the local environment to guide model building. Ramachandran and rotamer outliers were monitored and fixed as they appeared and then the chain was duplicated and rotated to fit a second interacting subunit density. The model was real-space refined using Phenix^59^ and repeated to create a model for three helical layers containing six peptide chains in total before a final real-space refinement in Phenix with NCS and template restraints applied to limit chain divergence. The final model quality was assessed using MolProbity^60^ and the results detailed **in Extended Data Table 2**.

### Mass spectrometry of App^NL-G-F^ Aβ fibrils

Sarkosyl-extracted fibrils were resuspended in 500 μl hexafluoroisopropanol (Sigma-Aldrich) and sonicated in an ultrasonic water bath (Untrawave Ltd. Cardiff) for 5 min. The mixture was incubated at room temperature for 24 hours and then centrifuged at 13,300 rpm for 5 min. The supernatant was filtered through a 0.22 μm PDVF filter (Millex-GV, Merck) and the filtrate was dried over a gentle stream of nitrogen gas to form a film of peptide around the wall of the tubes.

The sample was freeze-dried for 24 hours. The mixture was then resuspended in 50% acetonitrile aqueous solution with 2% formic acid and centrifuged at 13,300 rpm for 5 min. The sample was subsequently analysed by ESI-MS recorded using a Xevo QToF G2-XS mass spectrometer (Waters UK, Manchester, UK) operated in positive ion mode. Data were processed by using MassLynx V4.1 supplied with the mass spectrometer. The relative abundance of each peptide variant was calculated as relative abundance (%) = (ion count of peptide variant/total ion count of the five identified Aβ variants)X100%.

### Measurements of fibril width

12 tomograms in-tissue fibrils with the best contrast and a Methoxy-X04 cryoCLEM signal in CLEM were used for fibril width measurements. Four tomograms from *ex vivo* prepared amyloid were used to measure the fibril width of sarkosyl extracted Aβ. To measure width fibrils the outside edges were manually picked in IMOD of CTF corrected tomograms. The Euclidean distance between pairs of points was computed in Matlab. Welch t-test (α = 0.05) was performed using R to compare *ex-vivo* and in-tissue tomograms. One-way ANOVA and Tukey post-hoc test was performed to compare the distribution of fibril widths per tomogram.

### Data availability

The single-particle cryoEM refined maps, half-maps and model are deposited in the EMDB and PDB respectively as follows: the extracted *App*^*NL-G-F*^ fibril structure (16018/8BFA) and the control extracted Methoxy-X04-treated *App*^*NL-G-F*^ fibril structure (16019/8BFB). All tomographic datasets will be deposited at EMPIAR upon publication.

