## Extended Data Figures 1-10 for "The in-tissue molecular architecture of β-amyloid in the mammalian brain"

Extended Data Figure 1

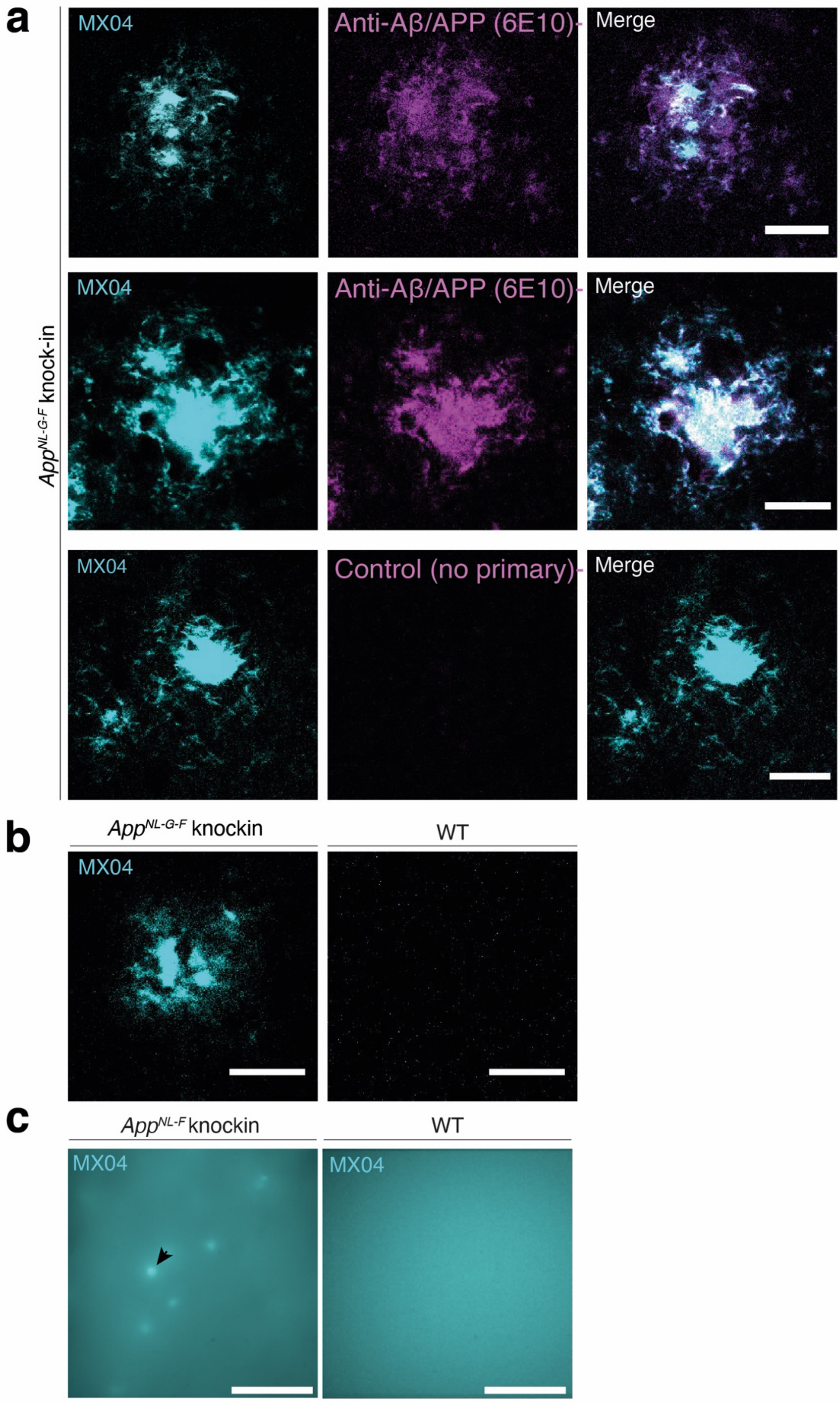


**Extended Data Figure 1. Immunohistochemical and cryoFM imaging of methoxy-X04 labelled amyloid plaques in *App$^{NL\text{-}G\text{-}F}$* mouse brain.**

(**a**) Immunofluorescence confocal microscopy images of amyloid plaques in fixed sections of *App$^{NL\text{-}G\text{-}F}$* mouse cortex. *Left panels*, Methoxy-X04-labelled plaques pseudo-coloured cyan.

*Middle panels*, APP/Aβ detected with 6E10 antibody pseudo-coloured magenta. *Right panels*, overlay of left and middle images.
*Bottom middle panel*, shows control sample in which 6E10 primary antibody was omitted. Scale bar, 50 µm.

(**b**) Immunofluorescence confocal microscopy images of fixed tissue sections stained with methoxy-X04 from $App^{NL\text{-}G\text{-}F}$ and wild-type control mouse cortex, *left* and *right* panels, respectively. Scale bar, 50 µm.

(**c**) Cryogenic fluorescence microscopy of high-pressure frozen mouse cortex from $App^{NL\text{-}G\text{-}F}$ and wild-type control mice that have received i.p. injection of methoxy X04 on *left* and *right*, respectively. Scale bar, 50 µm.

Extended Data Figure 2

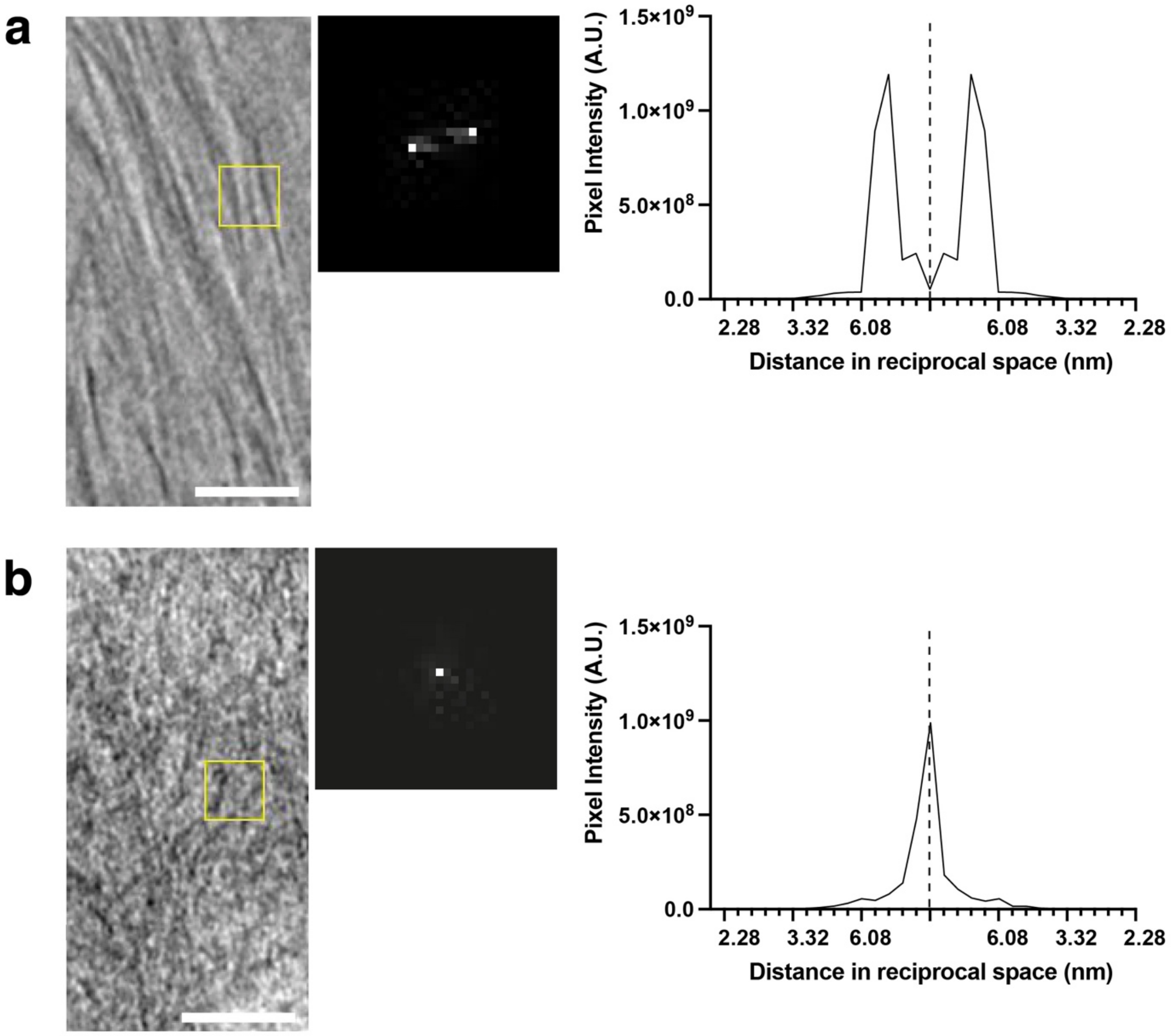


**Extended Data Figure 2**. **Quantitative assessment of the in-tissue architecture of amyloid plaques.**

(**a**) *Left*, Tomographic slice showing region with parallel bundles of amyloid from in-tissue tomograms of methoxy-X04 stained $App^{NL\text{-}G\text{-}F}$ amyloid plaque. Yellow box, region analysed by fast Fourier transformation (FFT) shown *Middle*. Scale bar, 20 nm. *Right*, profile of Fourier-transformed image showing reciprocal space distance versus pixel intensity (arbitrary units). A single peak away from the origin indicate the presence of parallel bundle. The peak indicates inter-fibril distance from the centre of one fibril to the next of amyloid plaques arranged in a parallel bundle.

(**b**) Same as **a** but for region showing amyloid not organised in a parallel bundle, indicated by a broad range of spatial frequencies.

Extended Data Figure 3

**a**

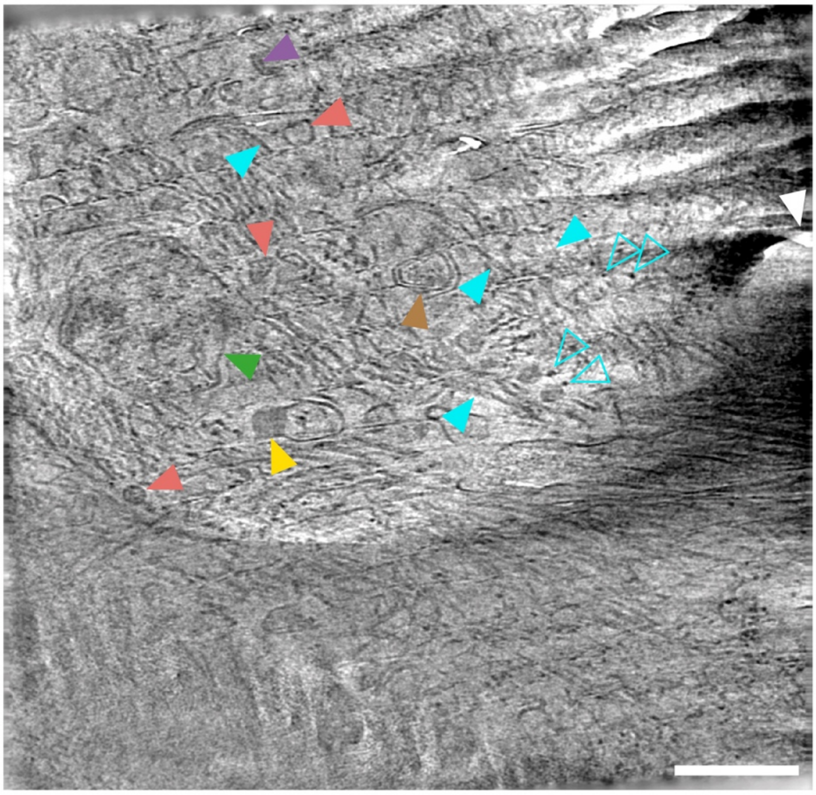

**b**

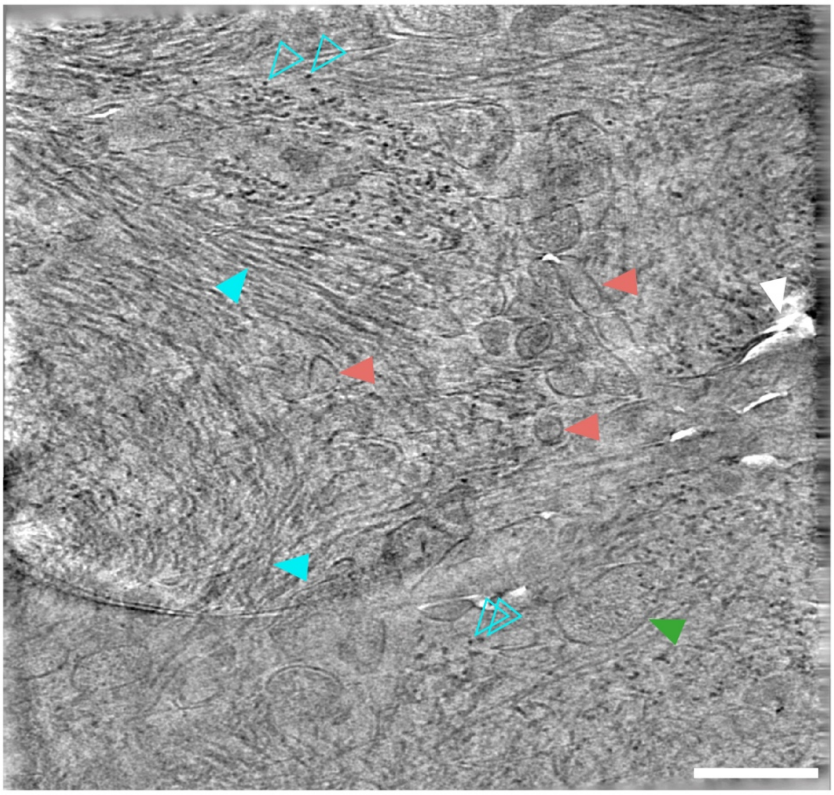

Extended Data Figure 3 continued

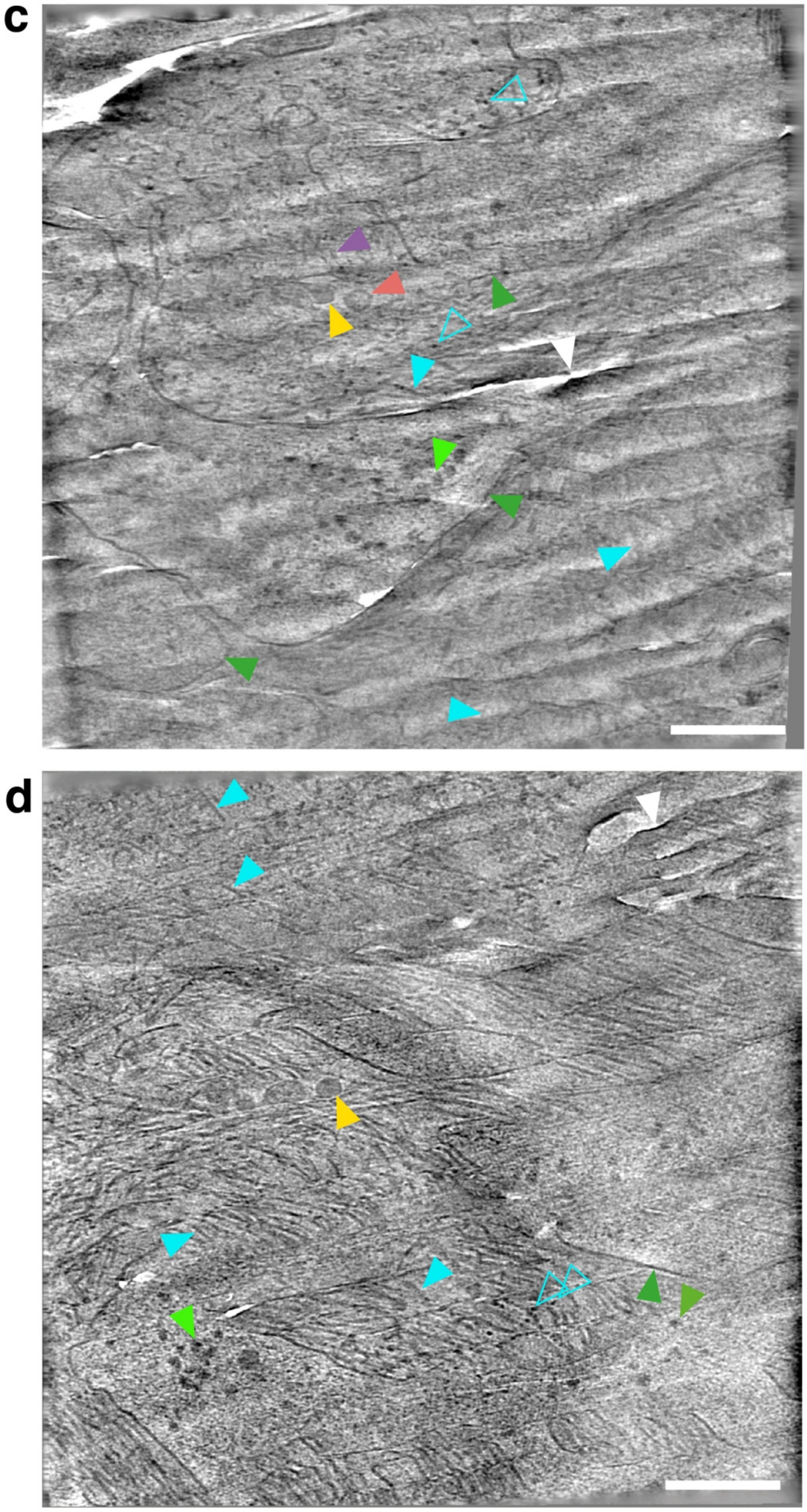


**Extended Data Figure 3**. **Representative tomographic slices from four tomograms. (a-d) collected within central regions of methoxy-X04-labelled amyloid plaques**. Scale bar, 100 nm. Filled and open cyan arrowhead, β-amyloid fibril oriented in the x/y plane and along the z-axis of the reconstructed tomogram, respectively. Red arrowhead, spherical exosome.

Purple arrowhead, squashed extracellular vesicle. Yellow arrowhead, extracellular droplet. Brown arrowhead, extracellular multilamellar body. Dark green arrowhead, subcellular compartment. Light green arrowhead, ribosome. White arrowhead, localised knife damage in tissue cryo-section. See methods section for criteria used to identify macromolecular and cellular constituents of tomograms. Scale bar, 100 nm.

Extended Data Figure 4

**a**

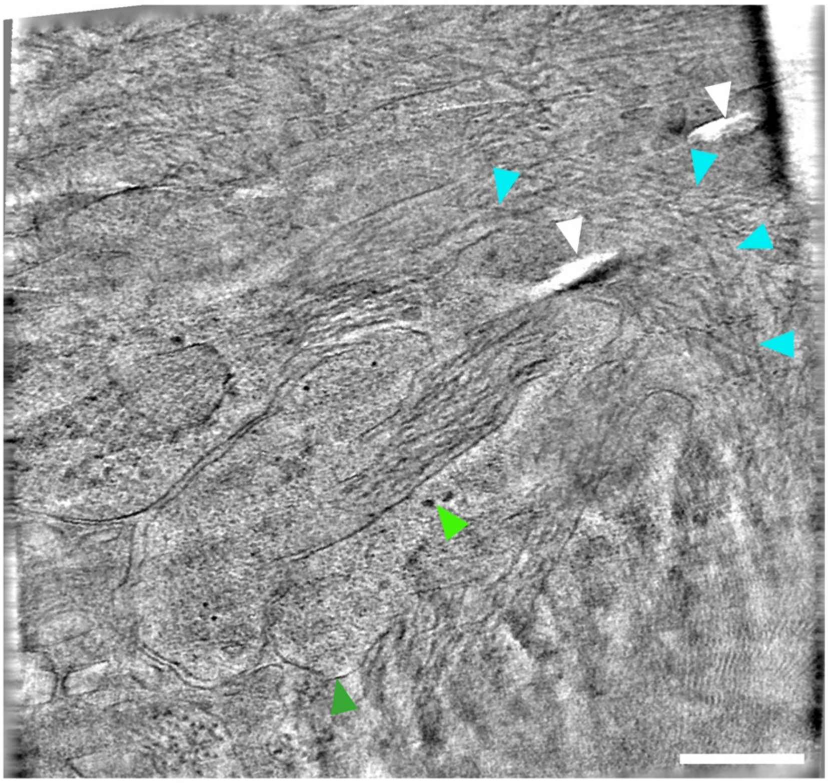

**b**

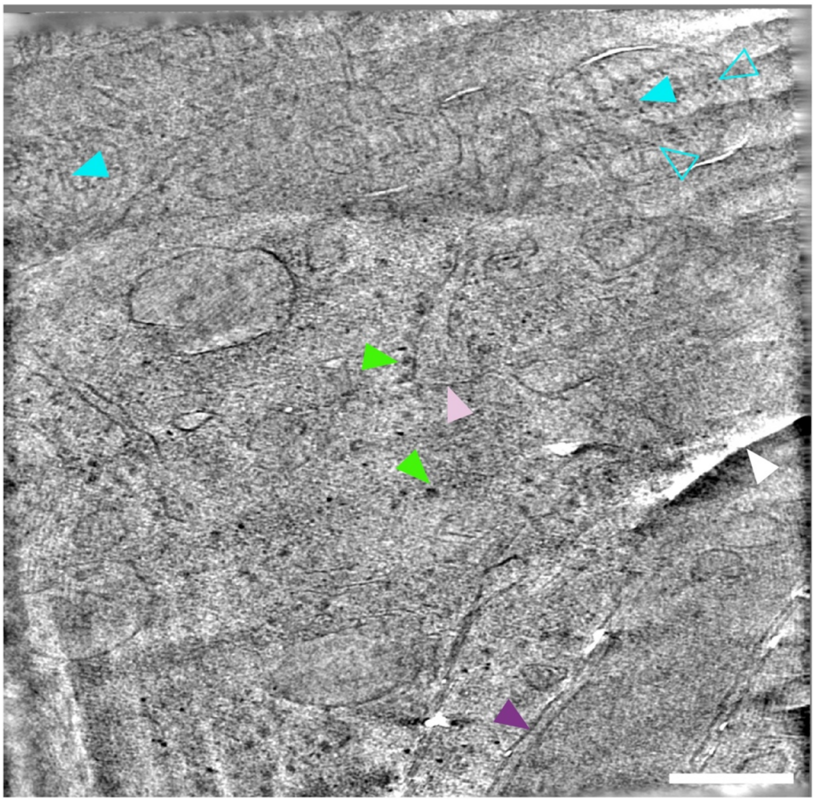

Extended Data Figure 4 continued

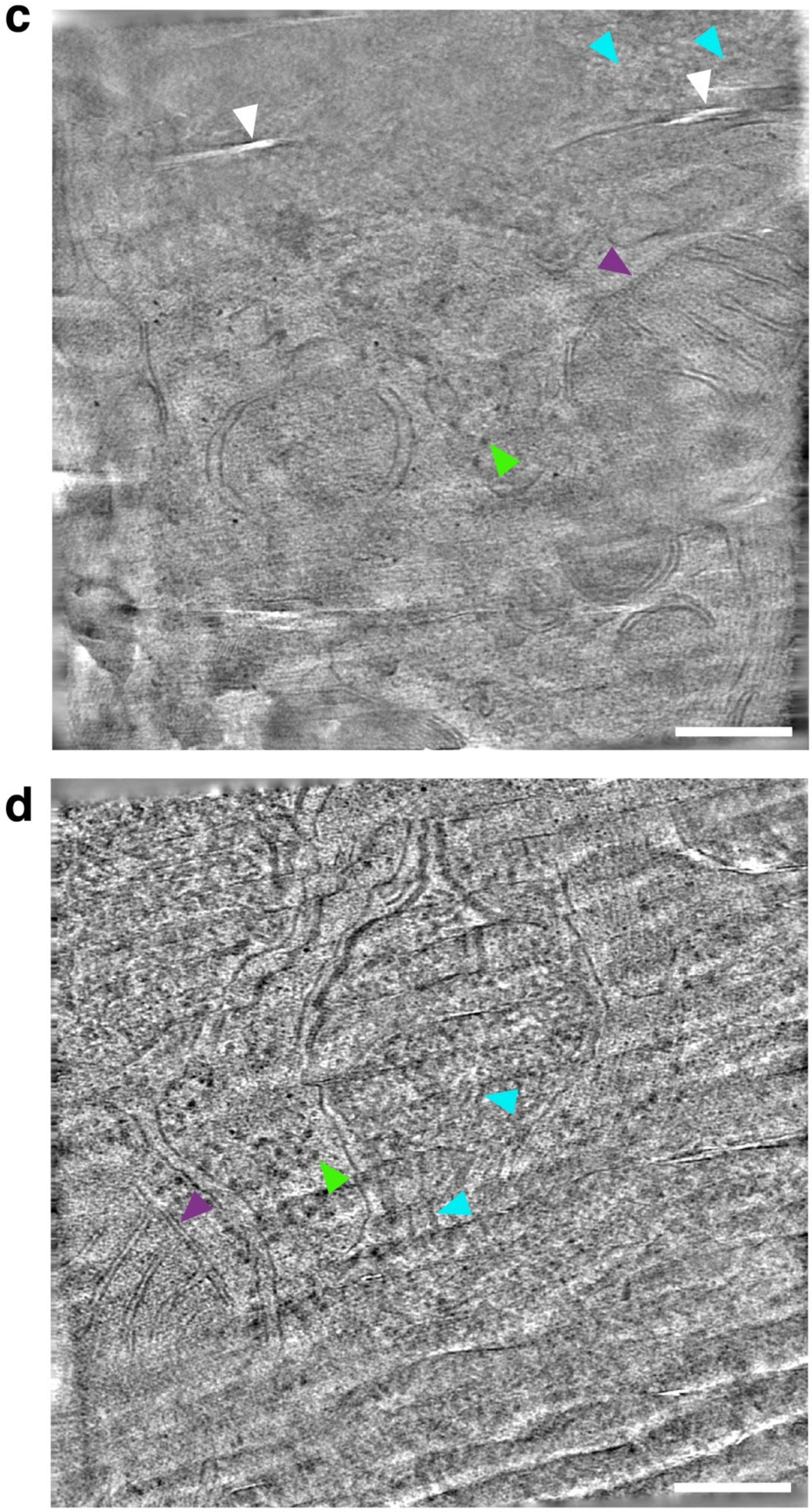


**Extended Data Figure 4**. **Representative tomographic slices from four tomograms (a-d) collected at peripheral regions of methoxy-X04-labelled amyloid plaques.** Scale bar, 100 nm. Filled and open cyan arrowhead, β-amyloid fibril oriented in the *x*/*y* plane and along the *z*-axis of the reconstructed tomogram, respectively. Purple arrowhead, mitochondria. Dark green arrowhead, subcellular compartment. Light green arrowhead, ribosome. White arrowhead, localised knife damage in tissue cryo-section. See methods section for criteria used to identify macromolecular and cellular constituents of tomograms. Scale bar, 100 nm.

Extended Data Figure 5

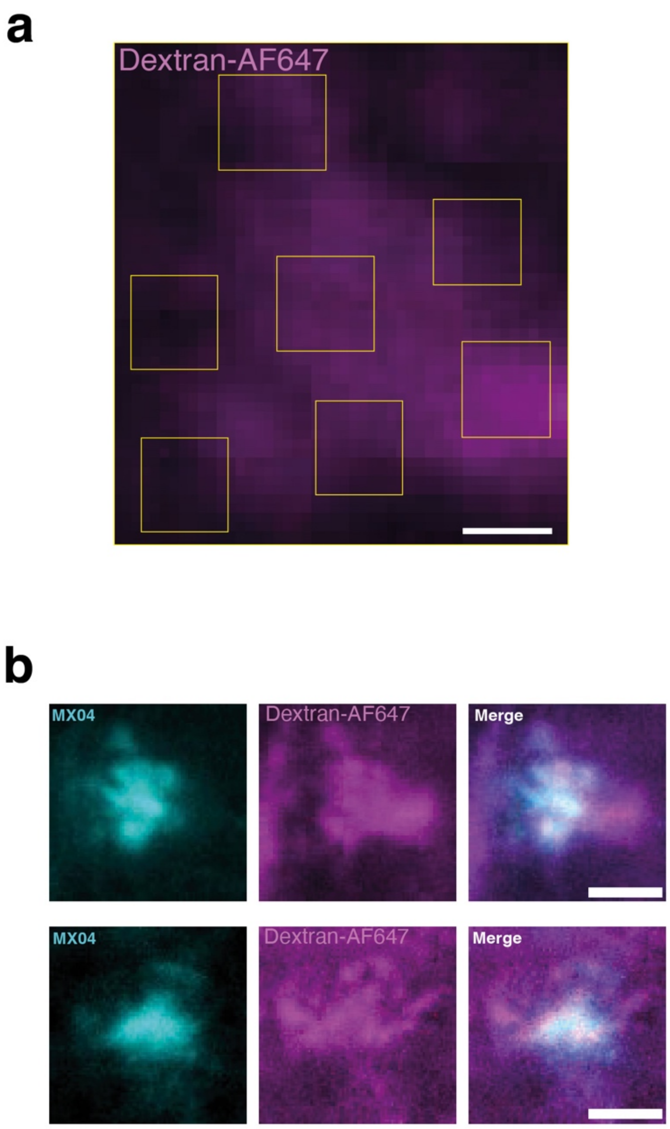


**Extended Data Figure 5**. **Cryogenic fluorescence microscopy of cryo-sections prepared from acute cortical slice of *App*$^{NL\text{-}G\text{-}F}$ mice that have received i.p. injection of methoxy X04.** Acute cortical slices were incubated with dextran-Alexa fluo-647 to label the extracellular space.

(**a**) Cryogenic fluorescence microscopy image of dextran-Alexa fluo-647-labelled tissue cryosections in magenta. Yellow boxes, indicate regions in which tomograms were collected (related to **Fig. 2c)**. Scale bar, 4 µm.

(**b**) Cryogenic fluorescence microscopy images of amyloid plaque within cryo-section. *Left*, methoxy-X04 pseudo coloured cyan. *Middle*, dextran-Alexa fluo-647 pseudo-coloured magenta. *Right*, overlay of left and middle images. Scale bar, 20 µm.

Extended Data Figure 6

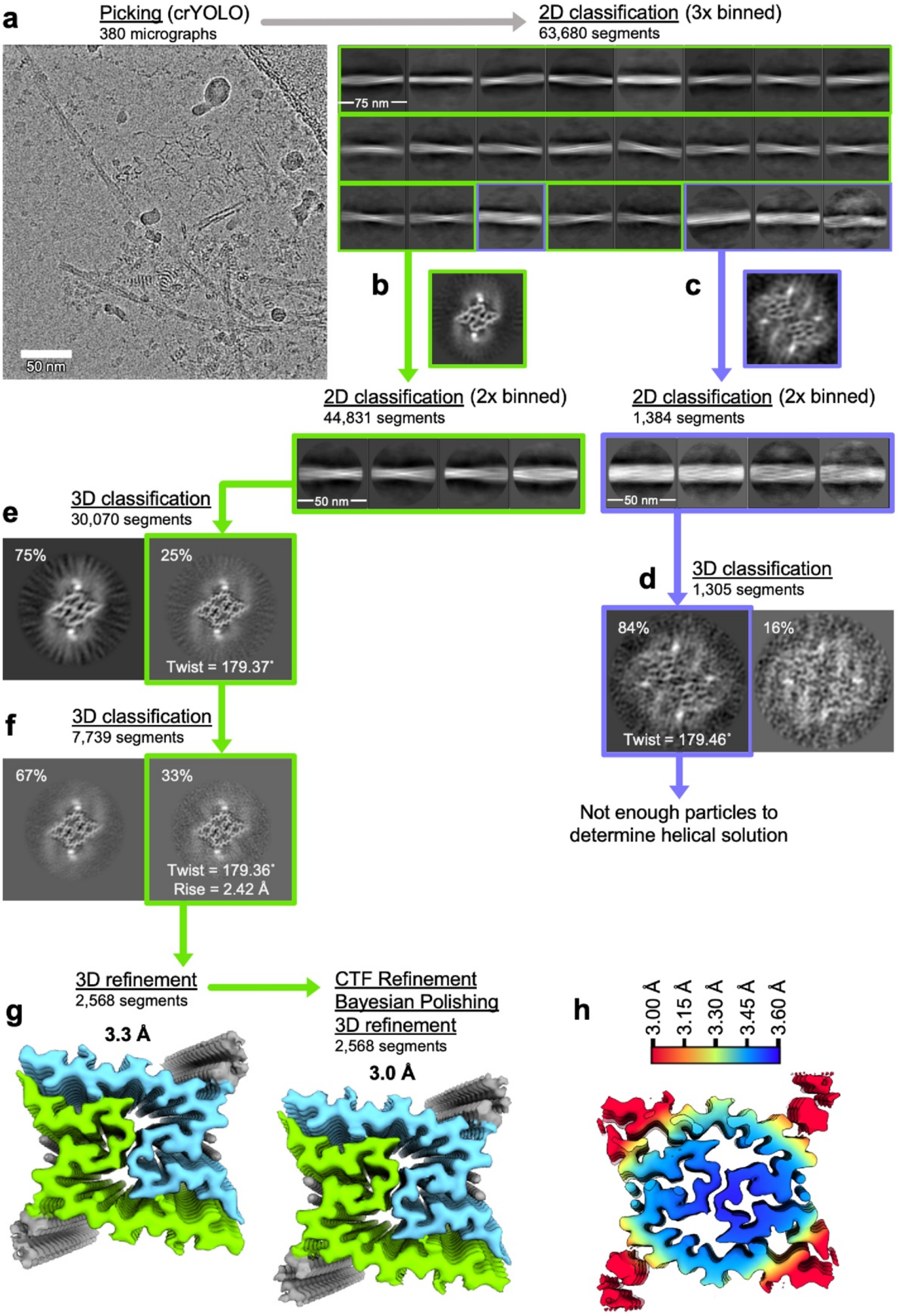


**Extended Data Figure 6. Single-particle cryoEM processing scheme for the extracted** ***App***$^{NL-G-F}$ **Aβ42 dataset.**

(**a**) A representative fibril-containing micrograph from the data, with scale bar length of 50 nm. The most populated 2D class averages are shown from the first round of classification of

the extracted helical segments. After removing picking artefacts, the data were split into two fibril subsets as highlighted by green or blue boxes.
(**b**) Representative 2D class averages from classification of the major fibril subset. Boxed inset shows an initial model for the form after 3D classification of a template generated from 2D class averages of the data.
(**c**) Representative 2D class averages from classification of the minor fibril subset. Boxed inset shows an initial model for the form after 3D classification of a template generated from 2D class averages of the data.
(**d**) Results of the first round of unbinned 3D classification of the minor fibril form, showing central slices of each output map. Despite further processing, the unambiguous fibril structure could not be determined.
(**e**) Results of the first round of unbinned 3D classification showing central slices of each output map which display the same polymorph but with different resolutions. The more ordered class was selected for further processing is highlighted in green.
(**f**) Results of the second round of unbinned 3D classification with helical searches, the selected output class is highlighted.
(**g**) The refined, sharpened maps before and after CTF refinement and polishing, each coloured by zone based on the two protein chains (as in **Fig. 4c**).
(**h**) Local resolution colouring of the final deposited map as calculated by RELION4.

Extended Data Figure 7

**a**

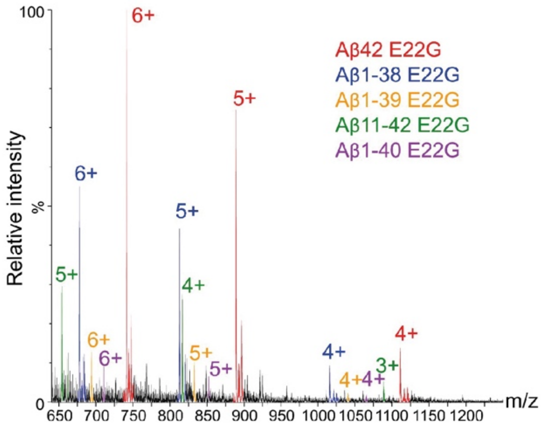


**c**

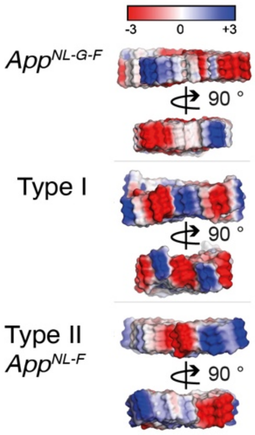


**b**

| Peptide identified | Sequence | Relative abundance | Theoretical MW (Da) | Observed MW (Da) | Charge state detected by ESI-MS | | | |
|---|---|---|---|---|---|---|---|---|
| | | | | | 6+ | 5+ | 4+ | 3+ |
| Ab42 E220G | DAEFRHDSGYEVHHQKLVFFA GDVGSNKGAIIGLMVGGVVIA | 51% | 4442.0 | 4441.2 | 741.203 | 889.246 | 1111.306 | ---- |
| Ab1-38 E22G | DAEFRHDSGYEVHHQKLVFFA GDVGSNKGAIIGLMVGG | 30% | 4059.5 | 4059.0 | 677.500 | 812.800 | 1015.735 | ---- |
| Ab11-42 E22G | EVHHQKLVFFAGDVGSNKGAII GLMVGGVVIA | 10% | 3063.8 | 3263.7 | ---- | 653.749 | 816.946 | 1088.576 |
| Ab1-39 E22G | DAEFRHDSGYEVHHQKLVFFA GDVGSNKGAIIGLMVGGV | 5% | 4158.6 | 4158.1 | 694.001 | 832.610 | 1040.518 | ---- |
| Ab1-40 E22G | DAEFRHDSGYEVHHQKLVFFA GDVGSNKGAIIGLMVGGVV | 4% | 4257.8 | 4257.1 | 710.516 | 852.420 | 1065.279 | ---- |

**d**

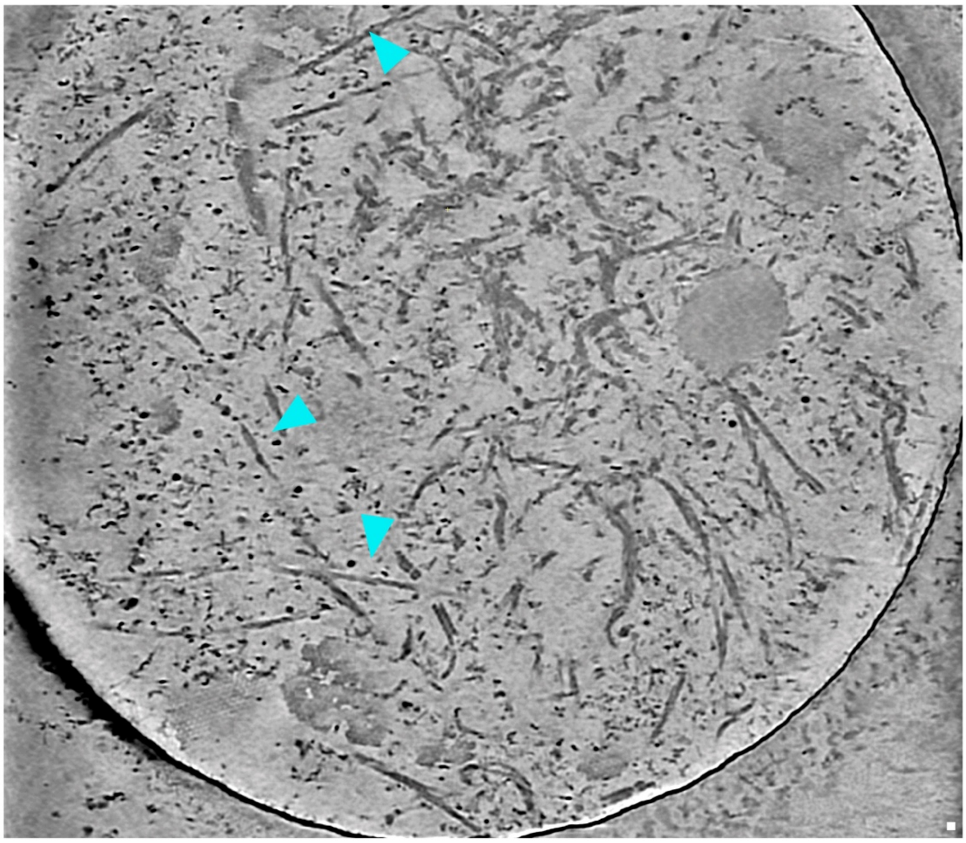

**Extended Data Figure 7**. **Mass spectrometry, solvent-exposed charge and cryoET of *ex vivo* β-amyloid.**

(**a**) Representative mass spectrometric detection of Aβ in sarkosyl-extracted ex vivo amyloid purified from *App*$^{NL\text{-}G\text{-}F}$ forebrain. Parent ions were identified as $A\beta_{1\text{-}42}$ E22G, $A\beta_{1\text{-}38}$ E22G, $A\beta_{11\text{-}42}$ E22G, and $A\beta_{1\text{-}40}$ E22G shown in red, blue, yellow, green and purple, respectively.
(**b**) Summary table showing, sequence, relative abundance, theoretical molecular weight, observed molecular weight, and charge states of Aβ peptide parent ions in sarkosyl-extracted *ex vivo* amyloid purified from *App*$^{NL\text{-}G\text{-}F}$ forebrain detected by mass spectrometry.
(**c**) Atomic models showing solvent-accessible surface coloured by charge of *App*$^{NL\text{-}G\text{-}F}$, Type I, and Type II Aβ fibrils.
(**d**) Representative tomographic slice through cryoET reconstruction of sarkosyl-extracted *ex vivo* amyloid purified from *App*$^{NL\text{-}G\text{-}F}$ forebrain. Cyan arrowheads, amyloid fibril exhibiting crossover points. Scale bar, 10 nm.

Extended Data Figure 8

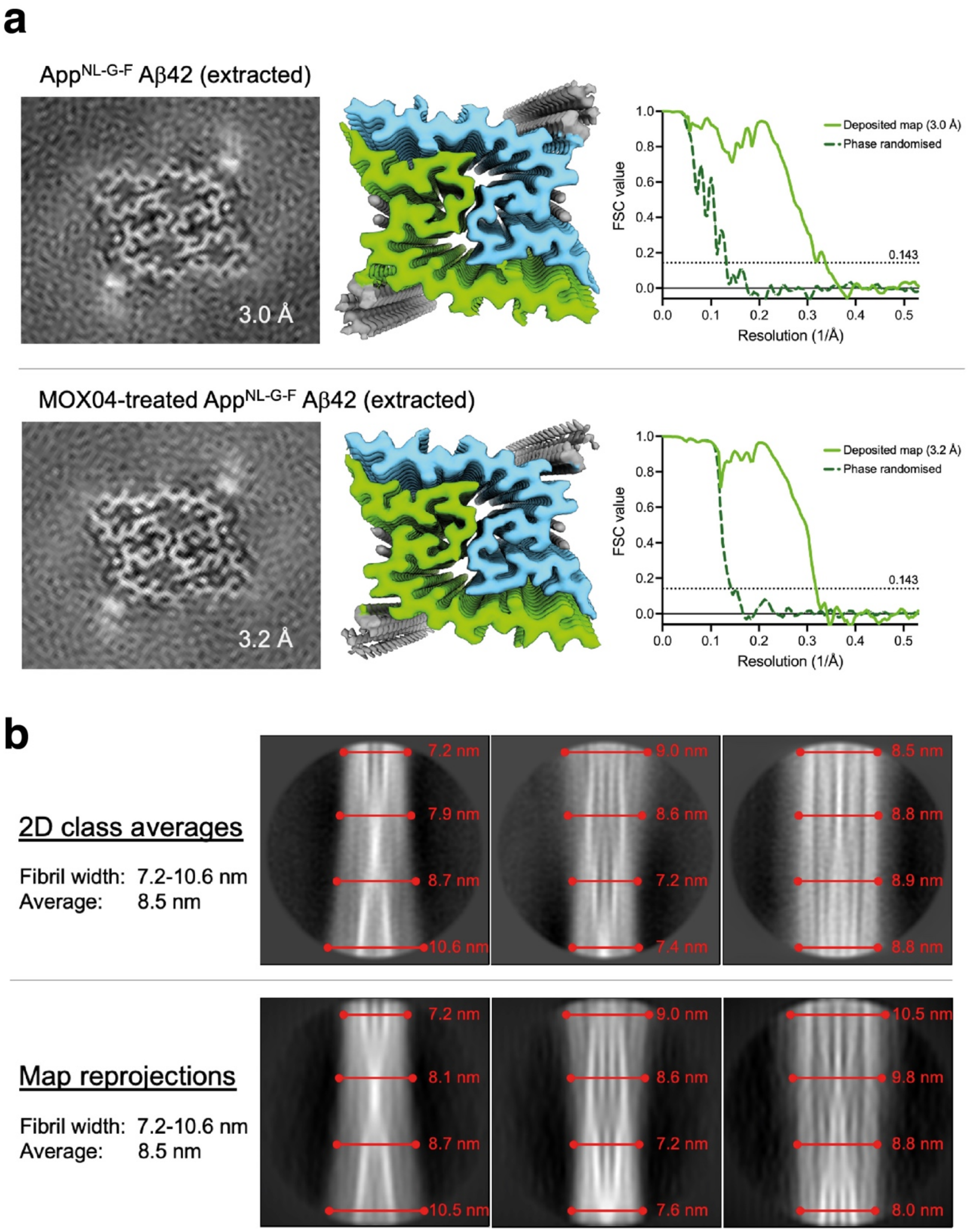


**Extended Data Figure 8. Cryo-EM of amyloid purified from MX04-labelled *App$^{NL-G-F}$* mice and fibril width measurements.**

(**a**) The deposited cryoEM maps from each extracted sample, from mice without (*above*) and with MOX04 staining (*below*). The central map slice shows a section corresponding to a single

helical layer from each, next to the cryoEM density map coloured by subunit and the FSC curves for the corrected masked map versus the phase randomised maps from postprocessing.
(**b**) Measurement of fibril widths from the extracted $App^{NL\text{-}G\text{-}F}$ single-particle cryoEM 2D class averages (*top*) and matched reprojections of the final map after lowpass-filtering to 10 Å (*lower*).

Extended Data Figure 9

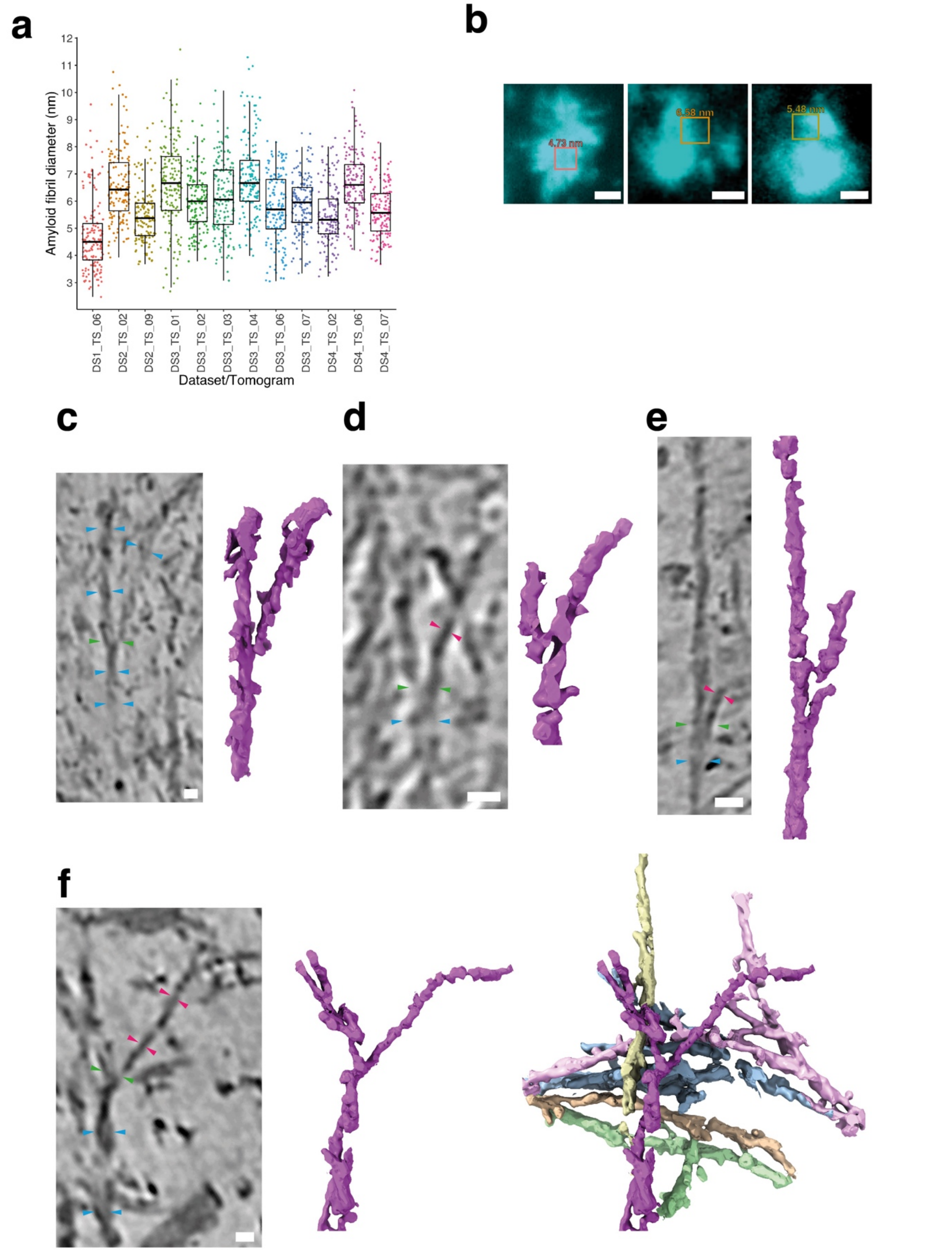


**Extended Data Figure 9. In-tissue and ex vivo cryoET evidence of protofilaments and branched amyloid.**

(**a**) Box plot showing distribution of fibril diameters (in nm) measured in 12 in-tissue tomograms of methoxy-X04 labelled amyloid in *App*$^{NL\text{-}G\text{-}F}$ mice.

(**b**) Cryogenic fluorescence microscopy images of methoxy-labelled amyloid plaueus *App*$^{NL\text{-}G\text{-}F}$ cryo-sections. Methoxy-X04 fluorescent signal is pseudo-coloured cyan. *Left*, amyloid plaque from which tomogram DS1_TS_06 was collected, highlighted by red box with an average fibril with of 4.73 nm. *Middle*, amyloid plaque from which tomogram DS2_TS_02 was collected, highlighted by red box with an average fibril width of 6.58 nm. *Right*, amyloid plaque from which tomogram DS2_TS_02 was collected, highlighted by light green box with an average fibril width of 5.48 nm. Scale bar, 10 µm.

(**c**-**e**) Examples of rare branched fibrils identified in cryoET data of sarkosyl-extracted ex vivo amyloid purified from *App*$^{NL\text{-}G\text{-}F}$ mice. *Left* panels, tomographic slices of ex vivo amylod cryoET reconstructions. Blue arrowheads, 4-13 nm fibril. Green arrowheads, fibril branch point. Magenta arrowheads, protofilament. Scale bar, 10 nm.

*Right* panels, raw tomographic density of branched amyloid.

(**f**) Same as (**c**-**e**) except *middle* panel shows raw tomographic density of branched fibril in magenta and *right* panels shows additional fibrils that contain branch points (in blue, pink, blue and green). Yellow and gold fibrils are single 3-5 nm diameter protofilaments.

Extended Data Figure 10

**a**

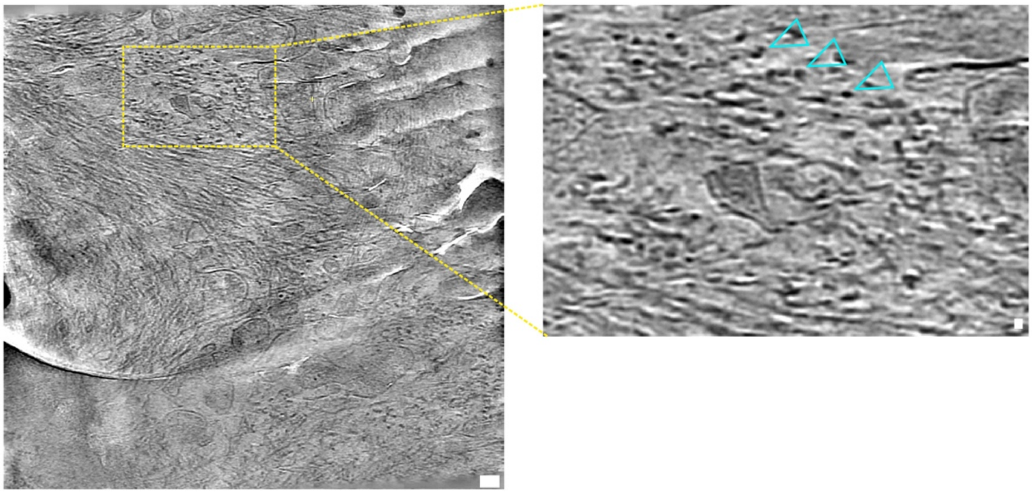

**b**

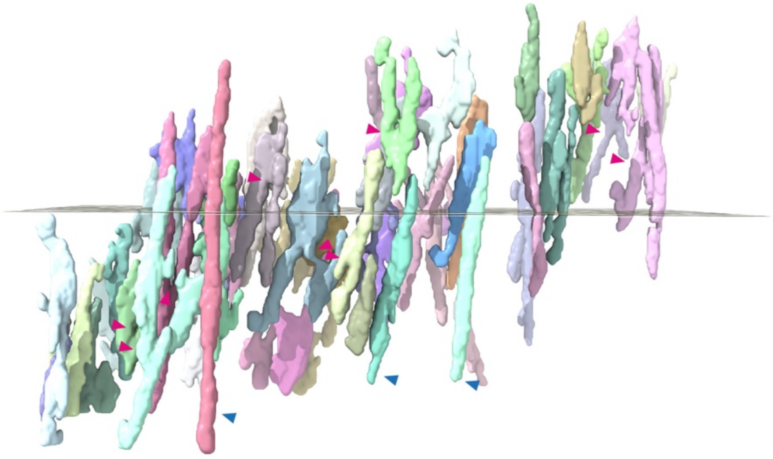

**Extended Data Figure 10. In-tissue tomogram showing additional examples of branched Aβ fibrils in *App*$^{NL\text{-}G\text{-}F}$ amyloid plaques.**

(**a**) *Left*, Tomographic slice of in-tissue amyloid with fibrils oriented on the z-axis of reconstructed tomographic volume. Scale bar, 50 nm. Dashed yellow box indicates close-up shown on *Right*. Cyan arrowheads, high-contrast spot corresponding to a single fibril. Scale bar, 10 nm.

(**b**) Segmented tomographic density of fibrils shown in (**b**), viewed from the side. Magenta arrowheads, putative branch points. Blue arrowheads, unbranched fibril.
